# A global-scale dataset of bat viral detection suggests that pregnancy reduces viral shedding

**DOI:** 10.1101/2024.02.25.581969

**Authors:** Evan A. Eskew, Kevin J. Olival, Jonna A. K. Mazet, Peter Daszak, PREDICT Consortium

## Abstract

Understanding viral infection dynamics in wildlife hosts can help forecast zoonotic pathogen spillover and human disease risk. Bats are particularly important reservoirs of zoonotic viruses, including some of major public health concern such as Nipah virus, Hendra virus, and SARS-related coronaviruses. Previous work has suggested that metapopulation dynamics, seasonal reproductive patterns, and other bat life history characteristics might explain temporal variation in spillover of bat-associated viruses into people. Here, we analyze viral dynamics in free-ranging bat hosts, leveraging a multi-year, global-scale viral detection dataset that spans eight viral families and 96 bat species from 14 countries. We fit hierarchical Bayesian models that explicitly control for important sources of variation, including geographic region, specimen type, and testing protocols, while estimating the influence of reproductive status on viral detection in female bats. Our models revealed that late pregnancy had a negative effect on viral shedding across multiple data subsets, while lactation had a weaker influence that was inconsistent across data subsets. These results are unusual for mammalian hosts, but given recent findings that bats may have high individual viral loads and population-level prevalence due to dampening of antiviral immunity, we propose that it would be evolutionarily advantageous for pregnancy to either not further reduce immunity or actually increase the immune response, reducing viral load, shedding, and risk of fetal infection. This novel hypothesis would be valuable to test given its potential to help monitor, predict, and manage viral spillover risk from bats.

## INTRODUCTION

The emergence of Ebola [1, 2], SARS-CoV [3], SARS-CoV-2 [4], and other zoonotic viruses has highlighted the urgent need to understand the processes that result in their spillover from animals to humans [5–9]. Because the exposure pathways that drive spillover are partially shaped by the prevalence and intensity of infection within animal hosts [6], understanding natural viral dynamics in wildlife may improve our ability to forecast and manage zoonotic disease risk. Bats are particularly important wildlife reservoirs of zoonotic viruses, including Marburg and other filoviruses, Nipah, Hendra, SARS-CoV, MERS-CoV, and SARS-CoV-2 [4, 10–13]. Further, some bat-associated viruses, such as Nipah and Hendra, show seasonal patterns of infection in intermediate hosts and humans [13–15]. These findings imply there are predictable elements of spillover and suggest that studying viral infection and shedding in bats may be valuable in developing strategies to prevent human disease [16, 17].

Seasonality of bat-associated viral outbreaks may be linked to features of bat life history and reproductive ecology, such as metapopulation structure, migration, or synchronous birthing [12, 13, 18–20]. For example, some bat species form large maternity colonies which increase local population density and intraspecific contact rates, thus increasing the potential for pathogen transmission and providing an ideal context for researchers to conduct viral sampling [18, 21, 22]. Viral infection in juvenile bats following the birthing period has attracted special scrutiny because seasonal influx of susceptible juveniles could facilitate pathogen maintenance within bat populations [18, 21–26]. Conversely, less attention has been given to maternal infection across reproductive states. Previous work has reported increased seropositivity [27–29] and viral detection [23, 30] in pregnant and/or lactating female bats. However, such studies are often restricted to sampling a relatively small number of bat host species and typically consider only a single viral pathogen, limiting their generalizability. Our understanding of bat viral dynamics during the energetically demanding periods of pregnancy and lactation [31, 32] would be enhanced by analysis of a broader range of host species and pathogens.

Here, we use hierarchical Bayesian models to examine the effects of reproductive status on viral detection using a global-scale dataset from bats sampled across 14 countries. We analyze viral detection data from eight viral families across 96 bat species belonging to nine families. Leveraging this large dataset, we explicitly account for important sources of variation, such as geographic region, specimen type, and different viral assays, to identify broad relationships between reproductive state and viral shedding across the order Chiroptera.

## METHODS

### Sample Collection

The majority of viral data analyzed here were from biological samples collected and tested as part of the PREDICT-1 project from 2009-2014. PREDICT-1 collaborated with partners in more than 20 countries throughout Asia, Africa, and Latin America to conduct virus surveillance, operating with the primary goals of public health capacity strengthening and pathogen discovery [33]. Sampling focused on human-livestock-wildlife interfaces in emerging infectious disease hotspots that were thought to represent a high risk of viral spillover of public health relevance [34, 35]. Field teams wearing appropriate personal protective equipment and trained in field biosafety protocols captured, sampled, and released all animals under veterinary supervision, with guidance from relevant local authorities and an IACUC protocol from the University of California, Davis (protocol number 16048) [36]. Here, we only use data from sub-adult and adult female bats (Table S1, Figure S1).

At the time of sample collection, individual bats were identified to the lowest taxonomic specification possible and sexed. Pregnancy and lactation status (yes/no) data were collected for female bats by gentle palpation of the abdomen and observation of milk production, respectively, following standard protocols [36, 37]. Non-invasive oral swabs, fecal or rectal swabs, urine or urogenital swabs, and blood samples were collected for later viral testing. Tissues (e.g., liver and spleen) were collected in rare cases when hunted bats were sampled or animals required euthanasia. As most specimens analyzed in our study were swabs (oral, rectal, or urogenital) or directly sampled excreta (feces and urine) that are likely routes of viral exposure to humans and other hosts, we believe that our confirmed positive samples approximate the occurrence of bat viral shedding. For oral and rectal swabs, viral detection may not always indicate direct infection of hosts and could instead represent viral sequence derived from food items, but this scenario is likely to be rare. Samples were primarily collected into either NucliSens® Lysis Buffer (bioMérieux, Inc., Marcy-I’Étoile, France) or TRIzol [38, 39]. They were frozen in liquid nitrogen in the field and safely transferred to in-country and US-based laboratories for storage at -80°C until viral testing.

### Viral Testing

Broadly reactive consensus PCR (cPCR) was used to detect both known and novel viruses [40]. Consensus PCR assays were targeted at either viral genus or family (details given in Table S2), with multiple testing protocols sometimes employed for the same focal viral taxon (e.g., two different assays were used for coronavirus detection). All cPCR positive samples were confirmed via sequencing and assigned a viral taxonomic designation by the PREDICT global laboratory team [39, 41].

### Statistical Modeling

We used hierarchical Bayesian models to analyze the association between reproductive status and viral infection in female bats while controlling for multiple potential confounders that are inherent in a complex, multi-country surveillance dataset. Our outcome of interest, viral detection, was the result (positive or negative) of sequence-confirmed viral cPCR tests on field-collected bat specimens. We modeled the association between female reproductive status and viral detection using a binomial distribution with a logit link function (Figure S2).

We performed initial cleaning and filtration steps on viral testing data to ensure quality and consistency (Figure S1). For example, we used only data from wild bat samples and excluded data from animals sampled in other settings (e.g., wildlife markets) because the stress associated with captivity could alter viral dynamics. In addition, while multiple discovery approaches can generate valuable viral diversity data [40], we sought to reduce potential bias in discovery efficacy by working solely with cPCR data, excluding results from high-throughput sequencing, real-time PCR, or serology methods. We also excluded data on viral groups that never resulted in viral detection across our female bat data subset. While these data could have been incorporated into our modeling efforts, they were ultimately uninformative and would bias downwards our estimates of overall viral detection probability.

After filtration (per above), data remained heterogeneous, with samples collected across host species, time, space, and other important categories. We account for this remaining variability in our models using varying effects. First, our models included varying intercepts and slopes (for pregnancy and lactation effects) for each bat host species (Figure S2). This approach allowed us to explicitly account for the fact that baseline viral detection and reproductive effects on viral detection could vary across host species. Varying intercepts and slopes also help to provide generalizable inference since this model structure allows us to recover community-level estimates for the intercept, pregnancy effect, and lactation effect parameters. In essence, by fitting varying intercepts and slopes by host species, we make use of data from all bat species to gain insight on the bat community generally. In addition, we constructed models that incorporated varying intercepts for the following data clusters: year of sample collection (which ranged from 2006-2014 because of the inclusion of archived specimens collected prior to the start of the PREDICT-1 project in 2009; see also [41]), country of sample collection, specimen type, the viral testing protocol used, and the diagnostic laboratory conducting testing (Figure S2). Table 1 provides a justification for the inclusion of each varying intercepts cluster. All varying intercepts clusters were modeled using normal priors with a mean of zero. Standard deviation parameters for these clusters were modeled using an exponential prior with a rate of 1. Finally, we implemented our varying intercepts clusters using a non-centered parameterization strategy to improve sampling efficiency and reduce bias [42]. This implementation recovers parameter estimates identical to those that could be obtained using a centered parameterization but improves the geometry of the posterior distribution for certain types of MCMC sampling problems. In sum, by incorporating these varying intercepts structures, we control for important sources of variation in our data (Table 1), therefore making more robust general inferences about viral dynamics in female bats.

**Table 1.**
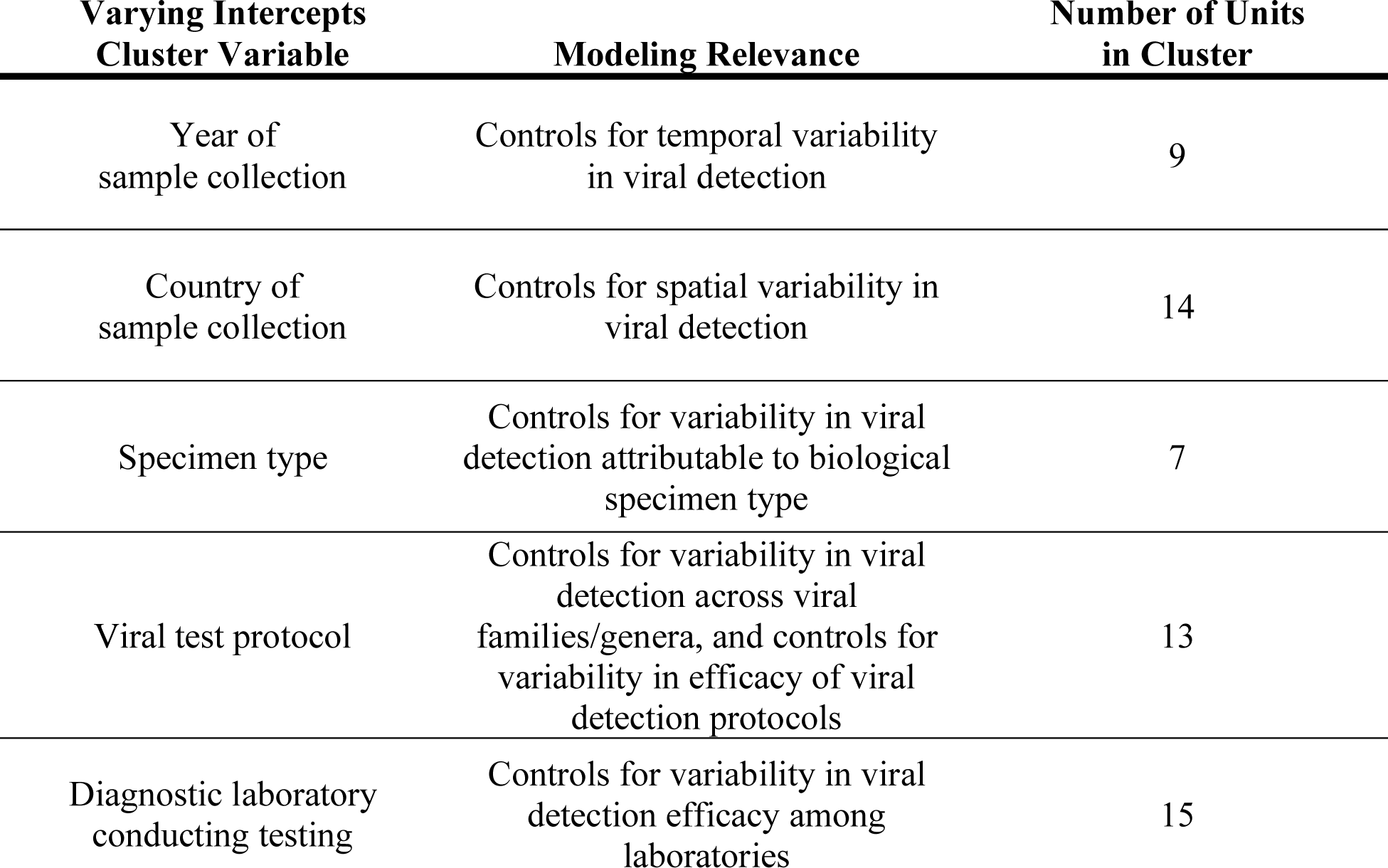
Variables included as varying intercepts clusters in hierarchical Bayesian models of viral detection in female bats. Variables are listed along with the justification for their inclusion. Within each varying intercepts cluster, individual units were fit with a unique intercept. In general, this strategy can be used to model mechanistic drivers of the outcome under investigation or as a way to control for “nuisance” variables. The numbers of distinct units within each varying intercepts cluster in the full female bat dataset are also shown. For technical details of the modeling approach, see the main text.

| Varying Intercepts Cluster Variable | Modeling Relevance | Number of Units in Cluster |
| --- | --- | --- |
| Year of sample collection | Controls for temporal variability in viral detection | 9 |
| Country of sample collection | Controls for spatial variability in viral detection | 14 |
| Specimen type | Controls for variability in viral detection attributable to biological specimen type | 7 |
| Viral test protocol | Controls for variability in viral detection across viral families/genera, and controls for variability in efficacy of viral detection protocols | 13 |
| Diagnostic laboratory conducting testing | Controls for variability in viral detection efficacy among laboratories | 15 |

Our varying intercepts structure for viral testing deserves special attention given its importance to our modeling results. Briefly, every cPCR test we analyzed is associated with both a general viral test type (i.e., the viral group that was tested for) and a specific viral testing protocol (i.e., the particular assay that was used). Using general viral test type as a control variable in our models would account for differences in detection across viral groups, but it would not account for differences in sensitivity or specificity associated with viral testing protocols. Therefore, our viral testing intercepts cluster consists of unique intercepts for each combination of general test type and specific testing protocol that was observed in the data (Table S2).

We chose to construct and fit a small suite of models that each contain the predictor variables that were considered relevant to our investigation *a priori* [43], namely the reproductive status variables. Conveniently, in a Bayesian setting, priors centered at zero have the same regularizing properties as shrinkage methods like ridge regression that are often used to control model complexity and reduce overfitting in large models [43–45]. This approach avoids some of the pitfalls associated with problematic ecological modeling strategies, including well-known issues with inference resulting from stepwise and all subsets model selection [44, 46–48]. Thus, we initially fit a model using the full female bat viral detection dataset (i.e., the *All Viral Families* dataset; Table S1, Figure S1). Then, to determine whether the effects of pregnancy and lactation on viral detection might differ depending on the viral taxa in question, we fit the model on data subsets representing testing from single viral families. We only fit single viral family data subsets when the subsets contained at least one viral detection from both pregnant and lactating bats to ensure relatively robust sample sizes across reproductive conditions. As a result, we fit additional models using data subsets for the following viral families: Adenoviridae, Coronaviridae, Herpesviridae, Paramyxoviridae, and Polyomaviridae. The Bayesian model we fit to the single viral family data subsets was identical to the model used to fit the *All Viral Families* dataset except in cases where varying effects clusters within the data subsets were only represented by one unit (i.e., only one level of a categorical variable). In those situations, we modified the model to exclude those varying intercept clusters in order to preserve model identifiability.

We specified and fit all models using the Stan programming language [49] through the ‘cmdstanr’ package interface [50]. We set Stan’s ‘adapt_delta’ and ‘step_size’ parameters to 0.99 and 0.5, respectively, to improve sampling of posteriors with difficult geometries at the expense of longer runtimes. For all model fits, we used four independent Markov chains, each with 3,500 iterations. Given that 1,000 iterations were used as warmup, we based our inferences on a total of 10,000 samples from each model (2,500 post-warmup iterations each from four chains). We performed post-hoc model diagnostics to ensure good model fits. These checks included visual inspection of chains using trace plots and confirmed convergence of the 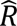 statistic towards one for all parameters [51]. We used the ‘cowplot’ [52], ‘dotwhisker’ [53], ‘ggplot2’ [54], ‘ggridges’ [55], and ‘rethinking’ [45, 56] packages to summarize and visualize model results. We report parameter estimates using posterior means and 95% highest posterior density intervals (HPDIs) to support model inference. In addition, where relevant, we draw attention to the proportion of a posterior’s probability mass that has support for values below or above zero (corresponding to probability of support for a negative or positive influence of the predictor on the outcome, respectively).

## RESULTS

### Data Summary

After data cleaning and filtering, our final female bat dataset contained 9,694 cPCR test results from 1,252 individuals of 96 different host species (Table S1, Figures S1 & S3). These data represent 459 detections of 123 unique viral species across eight viral families: Adenoviridae, Astroviridae, Coronaviridae, Herpesviridae, Paramyxoviridae, Parvoviridae, Polyomaviridae, and Rhabdoviridae.

In the female bat viral detection dataset, there were 28 bat host species for which we tested samples from both non-reproductive (not pregnant or lactating) and pregnant individuals. In 23/28 (82.1%) of those species, observed viral detection probability in samples from pregnant individuals was equal to or lower than that in non-reproductive individuals (Figure 1). There were 24 bat host species for which we tested samples from both non-reproductive and lactating individuals. In 21/24 (87.5%) of those species, observed viral detection probability in samples from lactating individuals was equal to or lower than viral detection in non-reproductive individuals (Figure 1).

**Figure 1.**
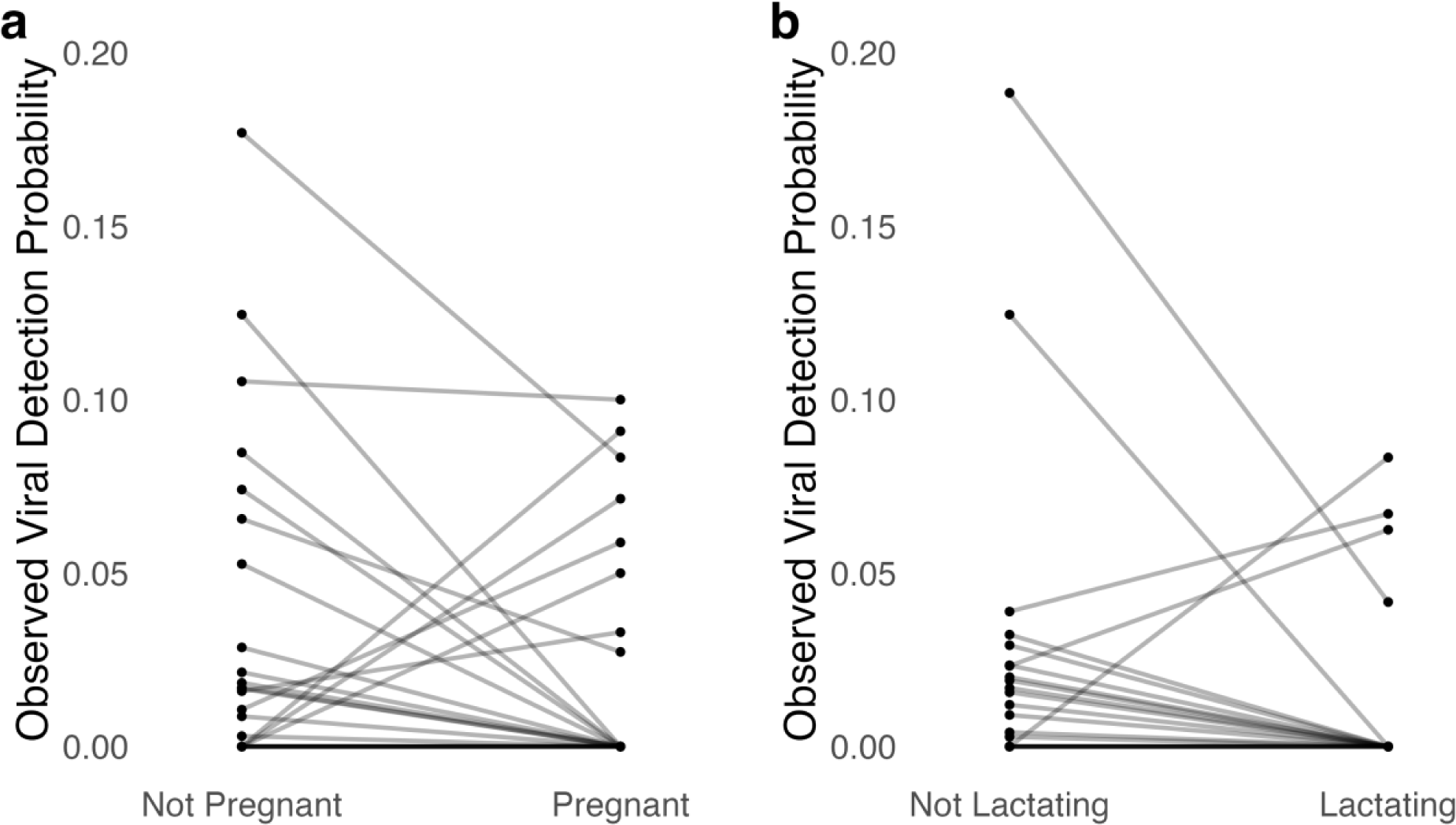
Observed effect of pregnancy (a) and lactation (b) on viral detection probability across bat host species. For visualization, the viral detection dataset was filtered to only include bat host species for which there were viral test data from both non-reproductive and reproductive conditions. All viral test data were pooled within host species and reproductive conditions. Lines connect data from the same host species and are displayed with transparency to emphasize areas of overlap in species trends.

### Hierarchical Bayesian Modeling

While the raw data summary suggests patterns of interest across reproductive categories, it does not account for important confounders in the surveillance dataset. Explicitly controlling for these factors motivated our use of hierarchical Bayesian models to investigate reproductive effects on viral detection in female bats. For all fit Bayesian models, the 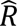 statistic for all parameters approached 1.0 (i.e., were < 1.01), and Markov chains mixed well (Figure S4). These diagnostics indicate model convergence. Further, we confirmed that the model fit to the full *All Viral Families* dataset was able to make accurate in-sample predictions: observed test positivity in the full dataset and for each viral family data subset fell within the 50% HPDI of model-based predictions (Figure S5).

In the full *All Viral Families* model, the posterior mean for the community-level intercept was negative (mean [95% HPDI]: -4.00 [-5.77, -2.26]), reflecting low viral detection overall (Figure 2). The 95% HPDI for the community-level pregnancy effect on viral detection spanned only negative values (-0.76 [-1.40, -0.02]; 97.4% posterior support for negative values), indicating strong support for a reduction in viral detection in pregnant female bats compared to non-gravid individuals (Figure 2). The community-level lactation effect’s posterior mean was also negative (-0.68), and although the 95% HPDI overlapped zero (-1.64, 0.19), 93.9% of the parameter posterior had support for negative values (Figure 2). In sum, these model estimates imply that modal viral detection probability in non-reproductive bats is ∼1%, while viral detection from pregnant and lactating individuals is roughly half that (∼0.4% detection probability; Figure 3). Estimates of species-specific reproductive effects closely conformed to the more general community-level estimates except in the case of heavily-sampled species such as *Eidolon helvum* and *Pteropus giganteus* (Figure S6).

**Figure 2.**
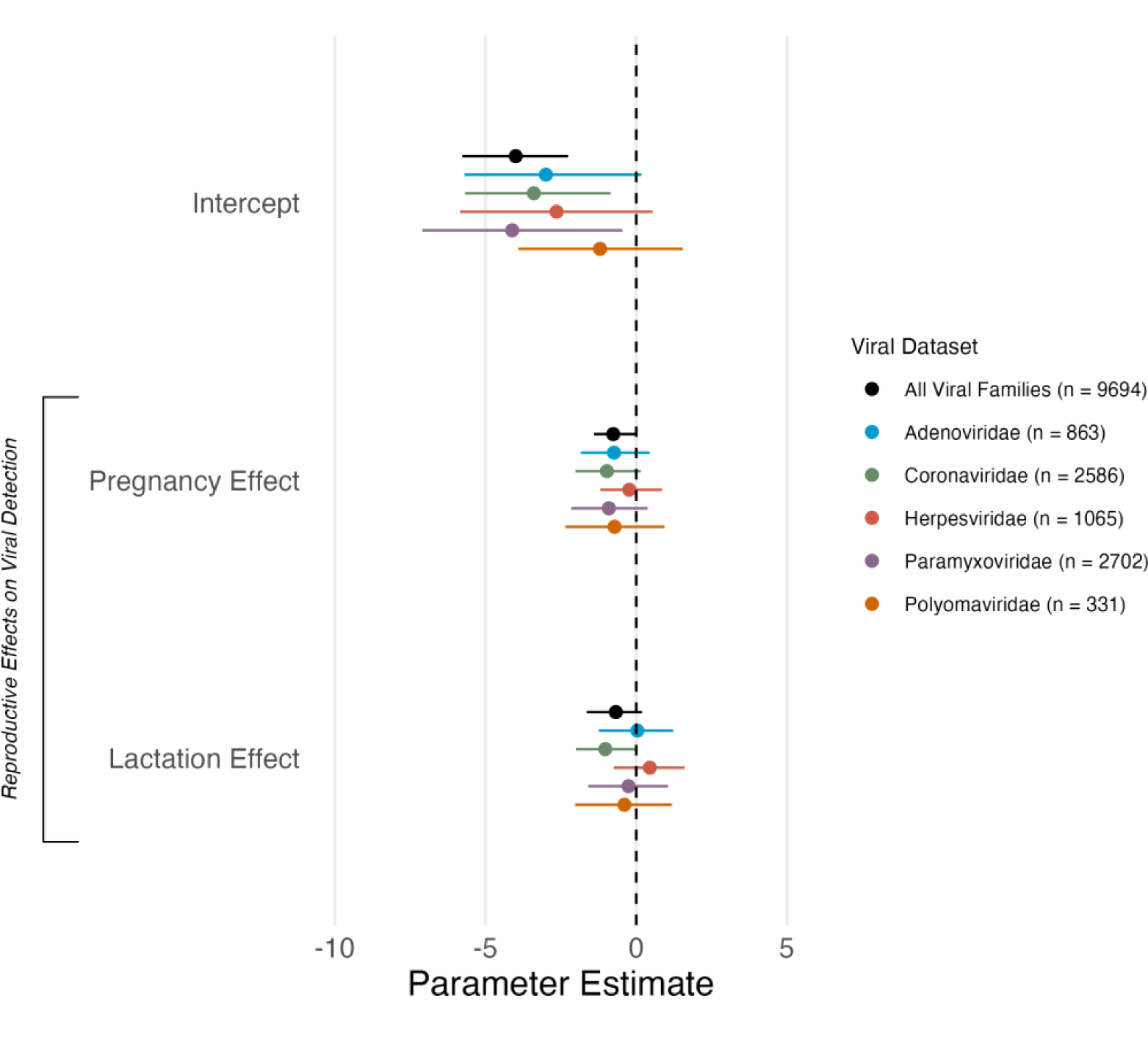
Dotchart showing parameter estimates from hierarchical Bayesian models of viral detection in female bats, including community-level intercepts, pregnancy effects, and lactation effects. Parameter means (dots) and 95% HPDIs are shown, and all parameter estimates are presented on the log-odds scale. The community-level intercept, pregnancy effect, and lactation effect parameters are shown for the full *All Viral Families* dataset as well as the five data subsets composed of data from single viral families. Sample sizes (number of cPCR tests) are indicated in the figure legend. Detailed description of the model structure, predictor variables, and model fitting is given in the main text and Figure S2.

**Figure 3.**
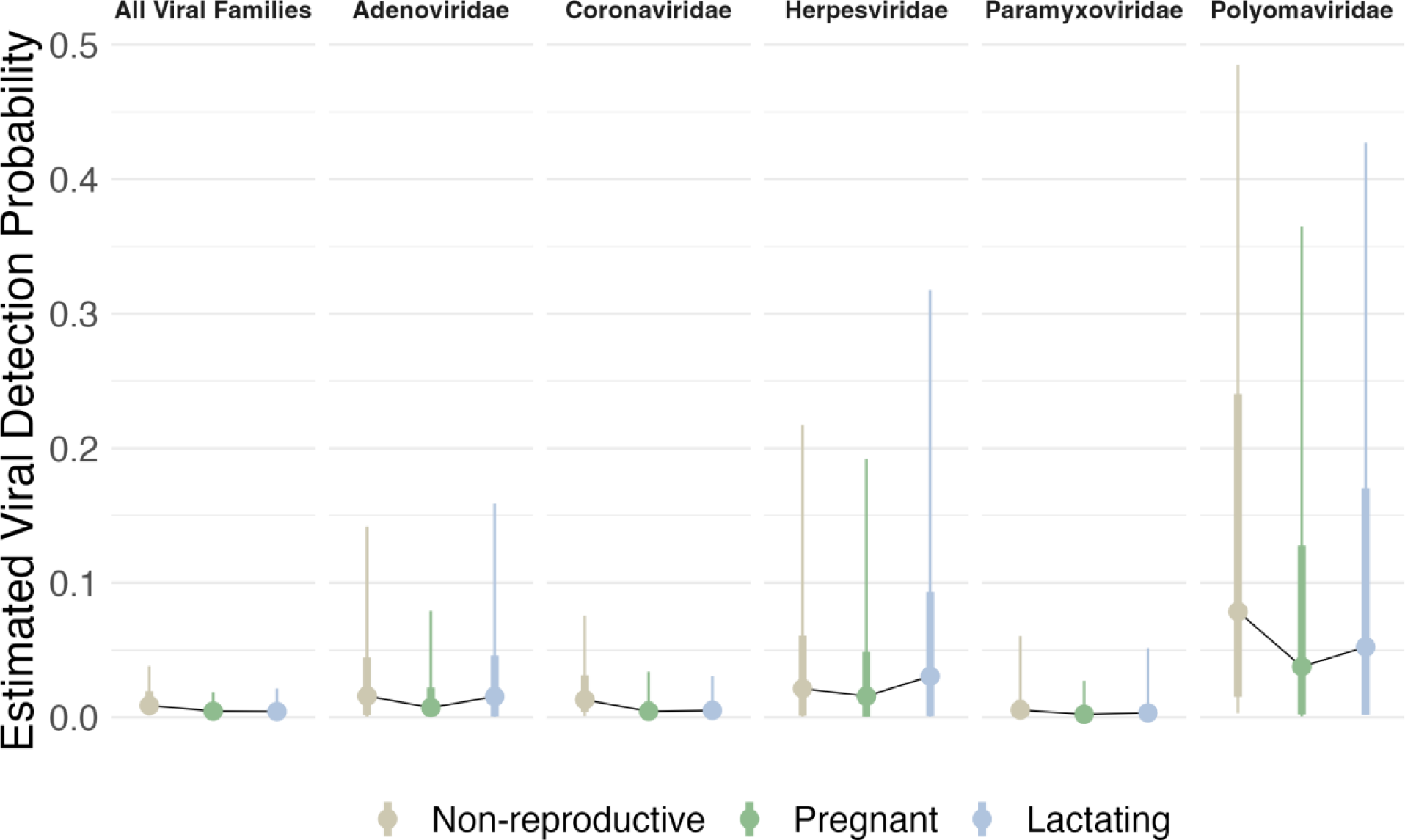
Model-based estimates of viral detection probability across reproductive conditions and viral datasets. Full parameter posterior distributions from fit models were used to generate estimates of detection probability for each reproductive condition-viral dataset combination. Points represent distribution modes, thin vertical lines represent 80% HPDIs, and thick vertical lines represent 50% HPDIs. Thin black lines chart the changes in modal viral detection probability across the non-reproductive, pregnant, and lactating conditions within each viral dataset. We elect to display modes rather than medians or means because for these highly right-skewed distributions, medians and means become unrepresentative of the location of the most concentrated probability mass.

Among the varying intercepts clusters in the model, the viral test protocol cluster had more variation than any other (Figure S7). This reflects the fact that the probability of viral detection varies strongly according to the viral family or genus being tested for and the protocol used (Table S2). Year and country of sample collection had more limited influence on viral test outcome, while specimen type and laboratory where analyses were conducted had the least effect of all (Figure S7). However, due to non-independent sampling across the varying intercepts cluster variables – for instance, different countries participating in different sampling years and samples from a country only being tested in specific laboratories – the individual effects of the varying intercepts are not easily interpretable despite being appropriately controlled for.

In examining parameter estimates from models fit to single viral family data subsets, the community-level intercepts for the Adenoviridae, Coronaviridae, Herpesviridae, and Paramyxoviridae data subsets were all similar to the estimate for the *All Viral Families* dataset (Figure 2). However, these estimates had inflated 95% HPDIs relative to the *All Viral Families* estimate, as would be expected given smaller sample sizes in the data subsets. In contrast, the community-level intercept for the Polyomaviridae data subset (-1.20 [-3.91, 1.55]) was noticeably higher than the *All Viral Families* estimate, indicating increased probability of viral detection in this group. In the single viral family data subsets, the community-level pregnancy effect posterior means were always negative (Figure 2). Although all estimates overlapped with zero in the 95% HPDI, the pregnancy effect posteriors for three of the data subsets (Adenoviridae, Coronaviridae, and Paramyxoviridae) had > 90% support for negative values (Figure 2). Thus, there was relatively consistent support across viral families for a negative effect of pregnancy on viral detection. Estimates of the community-level lactation effect from the single viral family subsets were more heterogenous. In three model sets (Adenoviridae, Paramyxoviridae, and Polyomaviridae), the 95% HPDI for the lactation effect parameter estimate substantially overlapped zero (Figure 2). The other two single viral family data subsets demonstrated opposing effects of lactation: in the Coronaviridae data subset, the community-level lactation effect posterior mean was negative (-1.03 [-2.00, -0.05], 98.4% posterior support for negative values), while in the Herpesviridae data subset, the effect was uncertain but likely positive (0.45 [-0.75, 1.61], 80.6% posterior support for positive values). Despite idiosyncrasies in the single viral family data subsets, overall our data implied lower viral detection in pregnant bats relative to non-reproductive individuals while detection in lactating bats tended to be similar to or somewhat lower than in non-reproductive bats (Figure 3).

## DISCUSSION

Our results highlight the important association between reproductive status and viral shedding in female bats. We consistently recovered a negative effect of pregnancy on viral detection in our global surveillance dataset after statistically accounting for other sources of variation inherent in this sampling design, including geographic and temporal heterogeneity. Lactation had a more uncertain influence on viral detection as the lactation effect differed in directionality across viral datasets. These findings have implications for our general understanding of viral dynamics during reproduction in chiropteran and mammalian hosts and can help inform zoonotic pathogen surveillance efforts that seek to sample wildlife during time periods of highest pathogen prevalence.

Pregnancy was the most important reproductive variable in our analyses, and model results across data subsets indicated that viral shedding is reduced in pregnant female bats. This finding contrasts with some previous studies that proposed pregnancy as a risk factor for viral infection in female bats [22, 27–29, 57]. However, these studies differ from ours in that they are of single bat populations or species and/or single viruses. Further, inference from these studies is sometimes complicated by relatively small sample sizes and/or use of serology, which indicates prior, but not necessarily active, infection [27–29, 57]. In contrast, our cPCR tests directly detect the presence of viral nucleic acid, indicative of infection and subsequent shedding. Furthermore, there are numerous reports of ambiguous associations between pregnancy and bat viral infection that also conflict with the prior studies that suggest a positive relationship between these two variables [23, 24, 30, 58].

Parasite and pathogen dynamics during bat reproduction are often contextualized using the idea of life history trade-offs, which posit that the costs of reproduction negatively impact other aspects of organismal performance, including host defense [59, 60]. Indeed, immune system depression is commonly invoked as an explanation when reproductive bats are found to have increased parasite prevalence or load [22, 26, 27, 61]. However, the costs of reproduction vary across taxa [62], such that immunological trade-offs during reproduction in any given focal taxon may not generalize. For example, the greater mouse-eared bat demonstrated depressed T cell response in early pregnancy [61], whereas the same immunological parameter in Brazilian free-tailed bats was not affected by reproductive status [63]. Similarly, in Daubenton’s bat, pregnant females had higher immunoglobulin G concentrations relative to non-reproductive individuals while lactating females had higher hemolysis titers, suggesting no obvious trade-off between reproduction and measures of immunity [64]. These disparities are perhaps not surprising given that findings in mammalian reproductive immunology often depend upon the host species and immunological components considered, as well as study methodologies [65–67]. In sum, the linkages between reproductive status, immune system function, and pathogen infection in bats are not currently well-resolved. Our study, which leveraged a standardized, global biosurveillance dataset to broadly address bat-virus interactions during reproduction, helps to clarify this subfield and serves as a platform for additional research to more critically approach this topic.

In humans, pregnancy was historically thought to drive immunosuppression in the mother and, consequently, increased susceptibility to infectious disease [68–70]. This paradigm has increasingly been challenged, however, and pregnancy may be most accurately described as a period of immune system modulation, one that does not necessarily imply broad immunodeficiency [68, 71]. For example, Kraus et al. [69] reported decreased activity of some immune cells, such as natural killer and T cells, during human pregnancy, but these shifts were accompanied by increased levels of defensin antimicrobial peptides and blood phagocytes. These findings can be reconciled with the fact that some diseases are known to be more severe during pregnancy [72, 73] if pregnancy induces immune system reorganization that simultaneously strengthens barriers to initial pathogen establishment while weakening immune control of existing infections [69, 70, 74]. Further, we now know of specific mechanisms, such as fetal-derived microRNAs, that can bolster maternal immune defenses during pregnancy [75]. This revised understanding of the immunology of pregnancy better matches evolutionary expectations, since there should be strong selective pressure for effective pathogen defense during the reproductive period as it is central to organismal fitness [68]. Collectively, these observations suggest that we should not assume that pregnancy strictly decreases immune defense and increases infection risk. Our statistical results, showing decreased viral detection in pregnant female bats, call for further integration of these perspectives into bat disease ecology in pursuit of a mechanistic understanding of wild bat immunology across reproductive states.

In contrast to strong pregnancy effects, we found more inconsistent evidence for an influence of lactation on bat viral detection. Although ∼94% of posterior probability mass supported a negative value for the lactation effect in our *All Viral Families* dataset, multiple datasets had estimates very close to zero, and mean effects within the Coronaviridae (negative) and Herpesviridae (positive) were contrasting. Lactation is costly and can demand more energy than pregnancy [31]. Thus, one might hypothesize that physiological trade-offs may be more severe during lactation than pregnancy, necessitating lower investment in immune defenses and viral control. Indeed, lactation status has been linked to increased viral detection in some bat systems [23, 30], but these results may be strongly confounded by reproductive behavior (i.e., lactating individuals in some species are often found in dense maternity roosts). On the other hand, although lactation does incur an energetic cost for reproductively active females, it should also represent a shift back towards normal physiological and immunological status following pregnancy. Dietrich et al. [22] reported decreased paramyxovirus prevalence immediately following parturition in *Mormopterus francoismoutoul*, a finding they attribute to the reestablishment of typical immune function in lactating females. Further, variation in the timing of sample collection within the lactation period (which differ in length across bat species) could have eroded any consistent signal of lactation on viral detection in our dataset. Regardless of the precise mechanisms at play, we find limited evidence that lactation reliably alters viral detection across the viral taxa considered, a result consistent with other recent work suggesting the likelihood of viral shedding during lactation is indistinguishable from non-reproductive periods [20].

Our findings point the way towards potentially fruitful avenues of investigation at the intersection of reproductive immunology and bat biology. In reproductive immunology, studies of humans and mice have begun to parse the complex immunological interactions between mother and fetus that contribute to a successful pregnancy, including fetal-derived antiviral factors that may influence maternal cells [75, 76]. While there are some similarities across taxa in immunological strategies during reproduction, the precise mechanisms that allow for maternal tolerance of the fetus while also providing for both maternal and fetal pathogen defense throughout gestation are likely species-specific [76]. Concurrent with these developments in reproductive immunology, a recent literature has begun to describe the unique immunological characteristics of bats, from their potential evolutionary origins [77–79] to their genetic basis [80–83]. These investigations have highlighted the ways in which bat immune systems are broadly distinct from other animals. However, our results, and the reproductive immunology work in other mammals cited above, suggest that detailed study of bat immune responses in the context of reproduction is warranted.

Bats appear to have immune responses that enable a level of tolerance to viral infection that is unique among mammals [79]. Field sampling and experimental infection data suggest bat species can harbor far higher viral loads at the individual-level, and far higher viral prevalence at the population-level, than other mammals. In the small number of bat species examined thus far, specific immunological differences appear to account for these patterns. These include constitutive interferon (IFN) activation and greater combinatorial diversity in immunoglobulin genes that do not undergo substantial affinity maturation [81]. In other studies, bats were shown to have dampened STING and suppressed inflammasome pathways, both of which contribute to immune tolerance and ultimately allow higher viral loads and prevalence [79]. We propose that it would be evolutionarily advantageous for pregnancy in bats to either not lead to further reduced antiviral activity or actually increase the immune response, so as to reduce viral load, shedding, and risk of fetal infection, concomitant with the findings in our study. This novel hypothesis would be valuable to test given its potential to help monitor, predict, and manage viral spillover risk from bats. Our findings also have relevance for understanding the role that vertical transmission plays in the maintenance of important zoonotic viruses in bats (e.g., Ebola) [84]. Experimental researchers should investigate the immune system remodeling that may occur near the fetal-maternal interface in bats, with special attention to the unique mechanisms that have been previously implicated in bat immune defense [79–83]. Studies that track immune defenses and viral exposure within cohorts of wild, reproductive female bats would also be especially valuable as a means to validate the statistical relationships and mechanistic hypotheses put forward here [85, 86].

Overall, our results improve our understanding of viral dynamics in bat hosts and can be used to inform viral surveillance and zoonotic disease prevention efforts. Our findings suggest that the intense focus on sampling maternity colonies in bat infectious disease research may be less efficient during pregnancy periods than previously assumed (given reduced viral detection during pregnancy) and could bias pathogen discovery. The description of new, potentially high-consequence viruses in bats outside of maternity roosts [38] emphasizes that temporally, geographically, and taxonomically broad sampling schemes will be key to comprehensive viral discovery and monitoring in bats [33]. Study designs incorporating multiple species and ecological contexts will also be necessary to completely characterize the viral diversity residing within mammalian host species, of which our knowledge remains incomplete [41, 87, 88]. Further, careful attention to viral dynamics throughout the reproductive cycle in a wider variety of bat host species will help identify periods of increased zoonotic spillover risk to humans, since spillover is often associated with pulses of viral excretion in reservoir hosts [6, 12, 24, 25]. Our work is congruent with recent, related studies that indicate the pup weaning period, not necessarily pregnancy or lactation periods, may be most strongly associated with viral shedding in bats [20, 89]. Consideration of the interplay between reproductive biology and host defenses will improve our understanding of virus-host dynamics and ultimately bolster our ability to prevent spillover from bats and other important wildlife hosts of viral pathogens.

## Acknowledgements

We thank the government partners whose generous collaboration enabled this research and the field teams and laboratory staff that contributed to sample collection and testing. Diego Montecino-Latorre, Nistara Randhawa, and Noam Ross gave feedback on early drafts that greatly improved this manuscript.

## Data Availability

Data and code supporting this manuscript are openly available via GitHub at https://github.com/ecohealthalliance/bat_viral_detection_reproduction.

