## Supplementary material for "A global-scale dataset of bat viral detection suggests that pregnancy reduces viral shedding": Figure S

**Table S1. Host species samples sizes in the full female bat viral detection dataset.** In each viral testing column, the first number represents the number of positive viral cPCR tests per host species, the second number represents the total number of cPCR tests per host species, and the proportion positive is given in parentheses. Host species (n = 96) are sorted alphabetically.

| **Host Species** | **All Viral Tests** | **Viral Tests from**  **Pregnant Individuals** | **Viral Tests from**  **Lactating Individuals** |
| --- | --- | --- | --- |
| *Acerodon celebensis* | 1 / 42 (0.02) | 0 / 0 (NA) | 0 / 0 (NA) |
| *Artibeus jamaicensis* | 0 / 164 (0) | 0 / 8 (0) | 0 / 17 (0) |
| *Artibeus lituratus* | 0 / 253 (0) | 0 / 30 (0) | 0 / 6 (0) |
| *Artibeus obscurus* | 1 / 31 (0.03) | 0 / 12 (0) | 0 / 0 (NA) |
| *Artibeus planirostris* | 1 / 76 (0.01) | 0 / 16 (0) | 0 / 12 (0) |
| *Balionycteris maculata* | 1 / 94 (0.01) | 1 / 11 (0.09) | 0 / 11 (0) |
| *Bauerus dubiaquercus* | 2 / 72 (0.03) | 0 / 2 (0) | 0 / 0 (NA) |
| *Carollia brevicauda* | 0 / 52 (0) | 0 / 3 (0) | 0 / 0 (NA) |
| *Carollia castanea* | 0 / 11 (0) | 0 / 0 (NA) | 0 / 0 (NA) |
| *Carollia perspicillata* | 6 / 279 (0.02) | 3 / 91 (0.03) | 0 / 23 (0) |
| *Carollia sowelli* | 5 / 70 (0.07) | 0 / 11 (0) | 0 / 0 (NA) |
| *Carollia subrufa* | 0 / 12 (0) | 0 / 0 (NA) | 0 / 0 (NA) |
| *Centurio senex* | 0 / 2 (0) | 0 / 0 (NA) | 0 / 0 (NA) |
| *Chaerephon pumilus* | 1 / 38 (0.03) | 1 / 20 (0.05) | 0 / 7 (0) |
| *Chiroderma trinitatum* | 0 / 1 (0) | 0 / 0 (NA) | 0 / 0 (NA) |
| *Choeroniscus godmani* | 0 / 2 (0) | 0 / 0 (NA) | 0 / 0 (NA) |
| *Choeroniscus minor* | 0 / 3 (0) | 0 / 3 (0) | 0 / 0 (NA) |
| *Coleura afra* | 3 / 64 (0.05) | 0 / 0 (NA) | 0 / 0 (NA) |
| *Cormura brevirostris* | 0 / 4 (0) | 0 / 0 (NA) | 0 / 0 (NA) |
| *Cynopterus brachyotis* | 0 / 60 (0) | 0 / 10 (0) | 0 / 0 (NA) |
| *Cynopterus horsfieldii* | 0 / 22 (0) | 0 / 0 (NA) | 0 / 0 (NA) |
| *Cynopterus sphinx* | 0 / 10 (0) | 0 / 0 (NA) | 0 / 10 (0) |
| *Dermanura gnoma* | 0 / 8 (0) | 0 / 0 (NA) | 0 / 8 (0) |
| *Dermanura phaeotis* | 0 / 50 (0) | 0 / 0 (NA) | 0 / 0 (NA) |
| *Dermanura watsoni* | 0 / 8 (0) | 0 / 0 (NA) | 0 / 2 (0) |
| *Desmodus rotundus* | 0 / 12 (0) | 0 / 0 (NA) | 0 / 0 (NA) |
| *Diphylla ecaudata* | 0 / 4 (0) | 0 / 4 (0) | 0 / 0 (NA) |
| *Dobsonia exoleta* | 1 / 13 (0.08) | 0 / 0 (NA) | 0 / 0 (NA) |
| *Eidolon helvum* | 90 / 515 (0.17) | 1 / 12 (0.08) | 2 / 48 (0.04) |
| *Epomophorus labiatus* | 2 / 240 (0.01) | 0 / 10 (0) | 0 / 20 (0) |
| *Glossophaga morenoi* | 0 / 5 (0) | 0 / 0 (NA) | 0 / 0 (NA) |
| *Glossophaga soricina* | 4 / 209 (0.02) | 0 / 22 (0) | 0 / 9 (0) |
| *Hipposideros caffer* | 5 / 41 (0.12) | 0 / 0 (NA) | 0 / 0 (NA) |
| *Hipposideros diadema* | 0 / 84 (0) | 0 / 0 (NA) | 0 / 0 (NA) |
| *Hipposideros galeritus* | 3 / 786 (0) | 0 / 0 (NA) | 0 / 52 (0) |
| *Hipposideros gigas* | 0 / 10 (0) | 0 / 0 (NA) | 0 / 0 (NA) |
| *Hipposideros larvatus* | 5 / 12 (0.42) | 0 / 0 (NA) | 0 / 0 (NA) |
| *Hipposideros ruber* | 1 / 26 (0.04) | 0 / 0 (NA) | 0 / 0 (NA) |
| *Hylonycteris underwoodi* | 0 / 6 (0) | 0 / 0 (NA) | 0 / 0 (NA) |
| *Kerivoula pellucida* | 0 / 11 (0) | 0 / 0 (NA) | 0 / 0 (NA) |
| *Lissonycteris angolensis* | 0 / 126 (0) | 0 / 0 (NA) | 0 / 10 (0) |
| *Micronycteris microtis* | 0 / 5 (0) | 0 / 0 (NA) | 0 / 0 (NA) |
| *Mimon crenulatum* | 0 / 20 (0) | 0 / 0 (NA) | 0 / 0 (NA) |
| *Miniopterus australis* | 0 / 11 (0) | 0 / 0 (NA) | 0 / 0 (NA) |
| *Miniopterus inflatus* | 6 / 95 (0.06) | 0 / 0 (NA) | 0 / 0 (NA) |
| *Miniopterus magnater* | 73 / 114 (0.64) | 0 / 0 (NA) | 0 / 0 (NA) |
| *Miniopterus schreibersii* | 0 / 13 (0) | 0 / 0 (NA) | 0 / 0 (NA) |
| *Molossus currentium* | 0 / 36 (0) | 0 / 36 (0) | 0 / 0 (NA) |
| *Molossus molossus* | 1 / 64 (0.02) | 0 / 5 (0) | 0 / 0 (NA) |
| *Murina suilla* | 0 / 11 (0) | 0 / 0 (NA) | 0 / 0 (NA) |
| *Myotis albescens* | 3 / 29 (0.1) | 1 / 10 (0.1) | 0 / 0 (NA) |
| *Myotis californicus* | 2 / 19 (0.11) | 0 / 0 (NA) | 0 / 0 (NA) |
| *Myotis nigricans* | 0 / 5 (0) | 0 / 0 (NA) | 0 / 4 (0) |
| *Myotis oxyotus* | 0 / 6 (0) | 0 / 0 (NA) | 0 / 0 (NA) |
| *Myotis velifer* | 0 / 3 (0) | 0 / 0 (NA) | 0 / 0 (NA) |
| *Nyctimene cephalotes* | 1 / 20 (0.05) | 0 / 0 (NA) | 0 / 0 (NA) |
| *Phoniscus atrox* | 0 / 10 (0) | 0 / 0 (NA) | 0 / 0 (NA) |
| *Phylloderma stenops* | 0 / 4 (0) | 0 / 0 (NA) | 0 / 4 (0) |
| *Phyllostomus discolor* | 0 / 22 (0) | 0 / 12 (0) | 0 / 0 (NA) |
| *Phyllostomus elongatus* | 1 / 12 (0.08) | 0 / 0 (NA) | 0 / 0 (NA) |
| *Phyllostomus hastatus* | 0 / 29 (0) | 0 / 21 (0) | 0 / 0 (NA) |
| *Pipistrellus coromandra* | 1 / 47 (0.02) | 0 / 0 (NA) | 1 / 47 (0.02) |
| *Platyrrhinus helleri* | 0 / 26 (0) | 0 / 0 (NA) | 0 / 7 (0) |
| *Pteronotus davyi* | 2 / 18 (0.11) | 0 / 0 (NA) | 0 / 0 (NA) |
| *Pteronotus parnellii* | 1 / 57 (0.02) | 1 / 14 (0.07) | 0 / 0 (NA) |
| *Pteropus alecto* | 14 / 507 (0.03) | 0 / 0 (NA) | 0 / 28 (0) |
| *Pteropus giganteus* | 149 / 3127 (0.05) | 40 / 1466 (0.03) | 65 / 969 (0.07) |
| *Pteropus lylei* | 0 / 16 (0) | 0 / 0 (NA) | 0 / 8 (0) |
| *Rhinolophus coelophyllus* | 6 / 13 (0.46) | 0 / 0 (NA) | 0 / 0 (NA) |
| *Rhinolophus creaghi* | 2 / 725 (0) | 0 / 60 (0) | 0 / 11 (0) |
| *Rhinolophus malayanus* | 0 / 6 (0) | 0 / 0 (NA) | 0 / 0 (NA) |
| *Rhinolophus paradoxolophus* | 1 / 41 (0.02) | 0 / 0 (NA) | 0 / 0 (NA) |
| *Rhinolophus sedulus* | 0 / 10 (0) | 0 / 0 (NA) | 0 / 0 (NA) |
| *Rhinolophus trifoliatus* | 0 / 50 (0) | 0 / 0 (NA) | 0 / 0 (NA) |
| *Rhinophylla pumilio* | 2 / 59 (0.03) | 0 / 0 (NA) | 1 / 16 (0.06) |
| *Rhynchonycteris naso* | 0 / 7 (0) | 0 / 0 (NA) | 0 / 0 (NA) |
| *Rousettus aegyptiacus* | 33 / 285 (0.12) | 0 / 20 (0) | 0 / 20 (0) |
| *Rousettus amplexicaudatus* | 6 / 91 (0.07) | 0 / 10 (0) | 0 / 0 (NA) |
| *Saccopteryx gymnura* | 0 / 4 (0) | 0 / 0 (NA) | 0 / 0 (NA) |
| *Scotophilus viridis* | 0 / 10 (0) | 0 / 0 (NA) | 0 / 0 (NA) |
| *Sturnira erythromos* | 0 / 6 (0) | 0 / 6 (0) | 0 / 0 (NA) |
| *Sturnira lilium* | 3 / 205 (0.01) | 1 / 17 (0.06) | 0 / 27 (0) |
| *Sturnira ludovici* | 2 / 111 (0.02) | 0 / 2 (0) | 0 / 6 (0) |
| *Sturnira magna* | 0 / 5 (0) | 0 / 0 (NA) | 0 / 0 (NA) |
| *Sturnira oporaphilum* | 0 / 6 (0) | 0 / 6 (0) | 0 / 0 (NA) |
| *Tadarida brasiliensis* | 1 / 5 (0.2) | 0 / 0 (NA) | 0 / 0 (NA) |
| *Taphozous mauritianus* | 1 / 36 (0.03) | 0 / 0 (NA) | 1 / 12 (0.08) |
| *Thoopterus nigrescens* | 0 / 50 (0) | 0 / 30 (0) | 0 / 0 (NA) |
| *Tonatia bidens* | 0 / 2 (0) | 0 / 0 (NA) | 0 / 0 (NA) |
| *Tonatia saurophila* | 1 / 6 (0.17) | 0 / 0 (NA) | 0 / 0 (NA) |
| *Trachops cirrhosus* | 0 / 10 (0) | 0 / 0 (NA) | 0 / 0 (NA) |
| *Triaenops persicus* | 13 / 59 (0.22) | 0 / 0 (NA) | 0 / 0 (NA) |
| *Trinycteris nicefori* | 0 / 10 (0) | 0 / 4 (0) | 0 / 0 (NA) |
| *Uroderma bilobatum* | 0 / 12 (0) | 0 / 7 (0) | 0 / 4 (0) |
| *Vampyriscus bidens* | 0 / 4 (0) | 0 / 0 (NA) | 0 / 0 (NA) |
| *Vampyrum spectrum* | 2 / 12 (0.17) | 0 / 0 (NA) | 0 / 0 (NA) |
| Total | 459 / 9694 (0.05) | 49 / 1991 (0.02) | 70 / 1398 (0.05) |

**Table S2. Viral test protocols included in the full female bat viral detection dataset.**

| **Viral Family** | **Target Viral Group** | **Target Gene** | **Reference** |
| --- | --- | --- | --- |
| Adenoviridae | Adenoviruses | Hexon | Casas, I., *et al*. 2005. Molecular identification of adenoviruses in clinical samples by analyzing a partial hexon genomic region. *Journal of Clinical Microbiology* 43: 6176-6182. |
| Adenoviridae | Adenoviruses | DNA polymerase | Wellehan, J.F., *et al.* 2004. Detection and analysis of six lizard adenoviruses by consensus primer PCR provides further evidence of a reptilian origin for the atadenoviruses. *Journal of Virology* 78: 13366-13369. |
| Astroviridae | Astroviruses | RNA-dependent RNA polymerase (RdRp) | Atkins, A., *et al*. 2009. Characterization of an outbreak of astroviral diarrhea in a group of cheetahs (*Acinonyx jubatus*). *Veterinary Microbiology* 136: 160-165. |
| Astroviridae | Astroviruses | RNA-dependent RNA polymerase (RdRp) | Chu, D.K.W., *et al*. 2008. Novel astroviruses in insectivorous bats. *Journal of Virology* 82: 9107-9114. |
| Coronaviridae | Coronaviruses | RNA-dependent RNA polymerase (RdRp) | Anthony, S.J., *et al*. 2015. Non-random patterns of viral diversity. *Nature Communications* 6: 8147.  *Modified from:*  Watanabe, S., *et al*. 2010. Bat coronaviruses and experimental infection of bats, the Philippines. *Emerging Infectious Diseases* 16: 1217-1223. |
| Coronaviridae | Coronaviruses | RNA-dependent RNA polymerase (RdRp) | Quan, P.-L., *et al*. 2010. Identification of a severe acute respiratory syndrome coronavirus-like virus in a leaf-nosed bat in Nigeria. *mBio* 1: e00208-10. |
| Herpesviridae | Herpesviruses | Terminase | Chmielewicz, B. *et al*. 2001. Detection and multigenic characterization of a novel gammaherpesvirus in goats. *Virus Research* 75: 87-94. |
| Herpesviridae | Herpesviruses | DNA polymerase | VanDevanter, D.R., *et al*. 1996. Detection and analysis of diverse herpesviral species by consensus primer PCR. *Journal of Clinical Microbiology* 34: 1666-1671. |
| Paramyxoviridae | Nipahviruses | N | Wacharapluesadee, S. and T. Hemachudha. 2007. Duplex nested RT-PCR for detection of Nipah virus RNA from urine specimens of bats. *Journal of Virological Methods* 141: 97-101. |
| Paramyxoviridae | Paramyxoviruses | RNA-dependent RNA polymerase (RdRp) | Tong, S., *et al*. 2008. Sensitive and broadly reactive reverse transcription-PCR assays to detect novel paramyxoviruses. *Journal of Clinical Microbiology* 46: 2652-2658. |
| Parvoviridae | Bocaviruses | NS1 | Kapoor, A., *et al*. 2010. Identification and characterization of a new bocavirus species in gorillas. *PLoS ONE* 5: e11948. |
| Polyomaviridae | Polyomaviruses | VP1 | Johne, R., *et al*. Novel polyomavirus detected in the feces of a chimpanzee by nested broad-spectrum PCR. *Journal of Virology* 79: 3883-3887. |
| Rhabdoviridae | Rhabdoviruses | RNA-dependent RNA polymerase (RdRp) | Anthony, S.J., *et al*. 2015. Non-random patterns of viral diversity. *Nature Communications* 6: 8147.  *Modified from:*  Bourhy, H., *et al*. 2005. Phylogenetic relationships among rhabdoviruses inferred using the L polymerase gene. *Journal of General Virology* 86: 2849-2858. |

**Figure S1. Flowchart of data cleaning and filtration steps used to generate the full female bat viral detection dataset.** Pink, rectangular compartments represent intermediate datasets, with the number of unique animals and unique species indicated in each. White, rounded compartments represent data filtration steps. Filtration criteria are indicated with R pseudocode and verbal descriptions. The green, rectangular compartment represents the full, *All Viral Families* viral detection dataset.

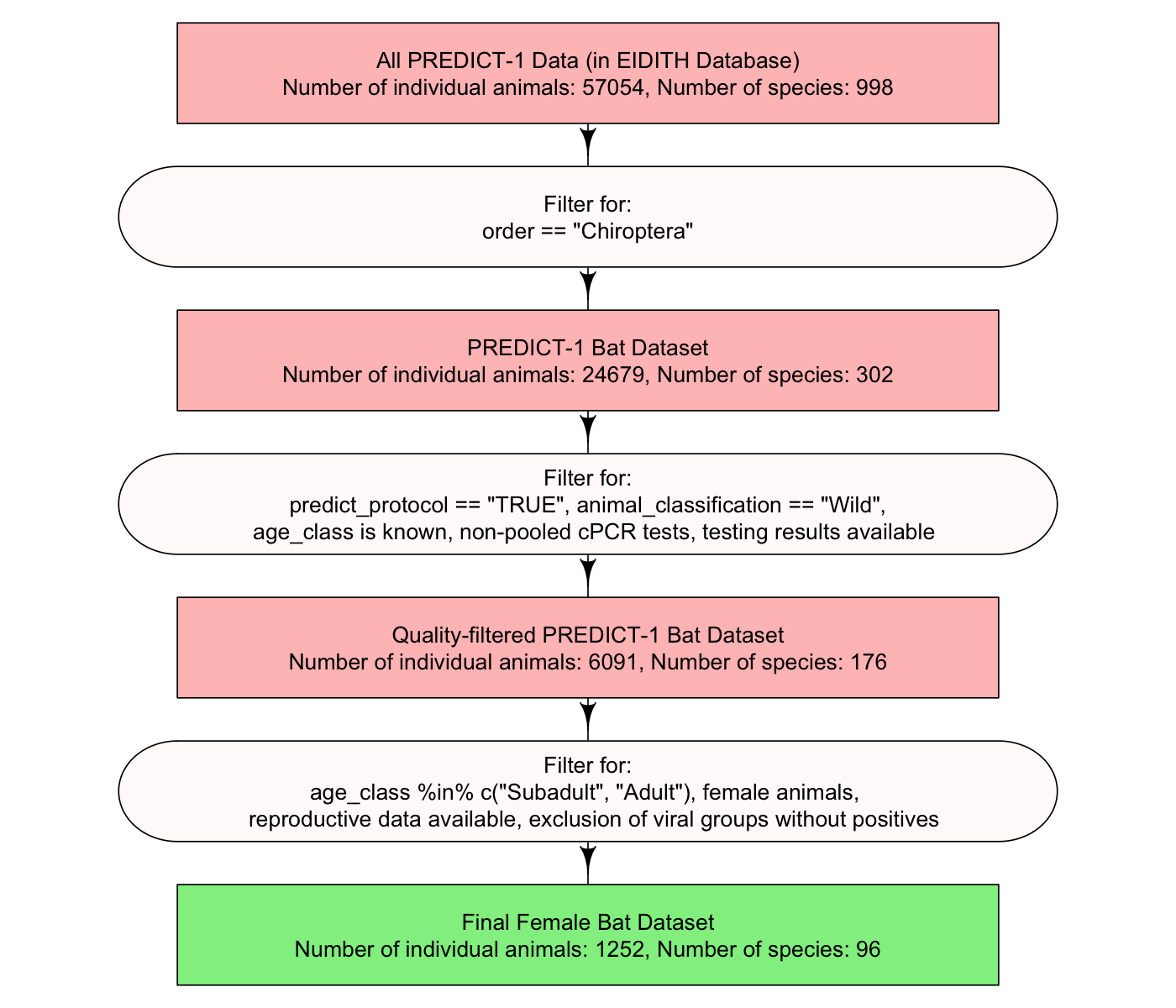

**Figure S2. Model structure for the hierarchical Bayesian model of viral detection in female bats.**

Likelihood

virus_detected*_i_* ~ Bernoulli(*p_i_*)

Linear model

logit(*p_i_*) ~ β_INTERCEPT,HOST SPECIES[_*_i_*_]_ +

β_PREGNANT,HOST SPECIES[_*_i_*_]_ * pregnant*_i_* +

β_LACTATING,HOST SPECIES[_*_i_*_]_ * lactating*_i_* +

α_YEAR[_*_i_*_]_ + α_COUNTRY[_*_i_*_]_ + α_SPECIMEN TYPE[_*_i_*_]_ + α_VIRAL TEST[_*_i_*_]_ + α_LABORATORY[_*_i_*_]_

Species-specific intercepts and reproductive effects drawn from multivariate normal distribution

$$\left[ \begin{matrix} \beta_{INTERCEPT,HOST SPECIES} \\ \beta_{PREGNANT, HOST SPECIES} \\ \beta_{LACTATING, HOST SPECIES} \end{matrix} \right] \sim MVNormal(\left[ \begin{matrix} \beta_{\mathrm{INTERCEPT}} \\ \beta_{\mathrm{PREGNANT}} \\ \beta_{\mathrm{LACTATING}} \end{matrix} \right], \Sigma)$$

$$\Sigma= \left( \begin{matrix} \sigma_{\beta_{\mathrm{INTERCEPT}}} & 0 & 0 \\ 0 & \sigma_{\beta_{\mathrm{PREGNANT}}} & 0 \\ 0 & 0 & \sigma_{\beta_{\mathrm{LACTATING}}} \end{matrix} \right) \Omega\left( \begin{matrix} \sigma_{\beta_{\mathrm{INTERCEPT}}} & 0 & 0 \\ 0 & \sigma_{\beta_{\mathrm{PREGNANT}}} & 0 \\ 0 & 0 & \sigma_{\beta_{\mathrm{LACTATING}}} \end{matrix} \right)$$

Priors for species-specific intercepts and reproductive effects

β_INTERCEPT_ ~ Normal(0, 2)

β_PREGNANT_ ~ Normal(0, 1)

β_LACTATING_ ~ Normal(0, 1)

$$\sigma_{\beta_{\mathrm{INTERCEPT}}} \sim Exponential(1)$$

$$\sigma_{\beta_{\mathrm{PREGNANT}}} \sim Exponential(1)$$

$$\sigma_{\beta_{\mathrm{LACTATING}}} \sim Exponential(1)$$

$$\Omega\sim LKJcorr(2)$$

Priors for varying intercept clusters

α_YEAR_ ~ Normal(0, σ_YEAR_)

α_COUNTRY_ ~ Normal(0, σ_COUNTRY_)

α_SPECIMEN TYPE_ ~ Normal(0, σ_SPECIMEN TYPE_)

α_VIRAL TEST_ ~ Normal(0, σ_VIRAL TEST_)

α_LABORATORY_ ~ Normal(0, σ_LABORATORY_)

σ_YEAR_ ~ Exponential(1)

σ_COUNTRY_ ~ Exponential(1)

σ_SPECIMEN TYPE_ ~ Exponential(1)

σ_VIRAL TEST_ ~ Exponential(1)

σ_LABORATORY_ ~ Exponential(1)

**Figure S3. Heat map of countries where female bats were sampled for viruses.** As indicated in the figure legend, intensity of red color corresponds to the number of cPCR tests in our full female bat viral detection dataset conducted on samples collected from each country.

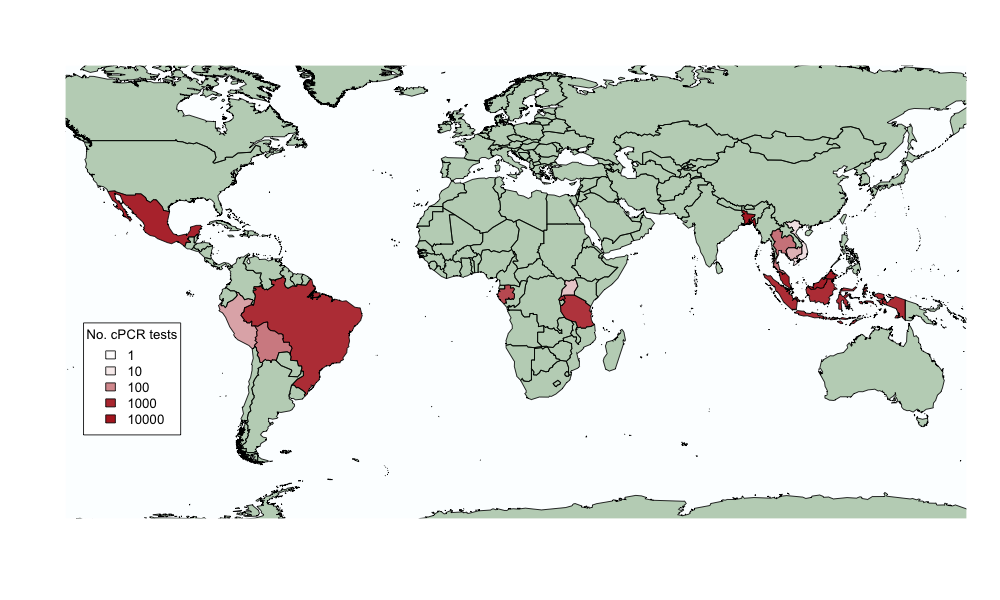

**Figure S4. Trace plots for select parameter estimates from a hierarchical Bayesian model of viral detection in female bats.** Plots show traces across all four independent Markov chains, with each chain represented in a different color. The first three parameters (the community-level intercept and reproductive effects) are identical to those shown for the *All Viral Families* dataset in main text Figure 2. Note that the y-axis scale differs across parameters.

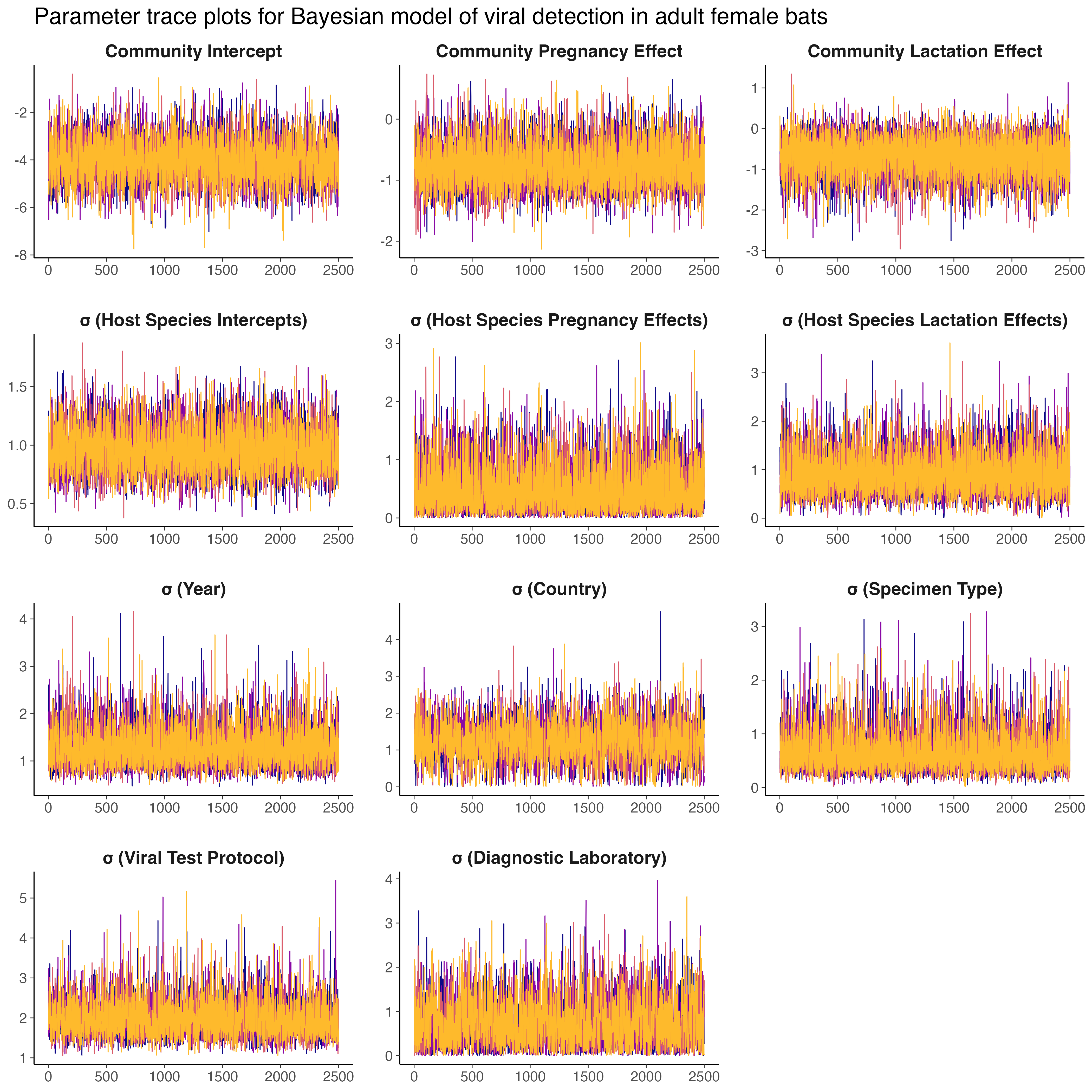

**Figure S5. In-sample predictions derived from the hierarchical Bayesian model fit to the full *All Viral Families* dataset.** To confirm that our model was able to adequately capture the observed data, we used posterior parameter values across all model iterations (n = 10,000) and covariate values associated with all of the observed viral detection data (n = 9,694) to generate model-based predictions of viral detection. For each set of posterior parameter values, these model-based predictions were summarized into a test positivity metric (predicted positive viral detections / 9,694 viral cPCR tests). Predicted test positivity was calculated across the full dataset and separately for each viral family targeted by testing (n = 8) to verify that the model performed well across this important dimension of the dataset. For each data subset, the thin grey lines represent the 95% HPDI of test positivity predictions while the thick grey box represents the 50% HPDI predictive interval. Observed test positivity is shown for each data subset as a red point.

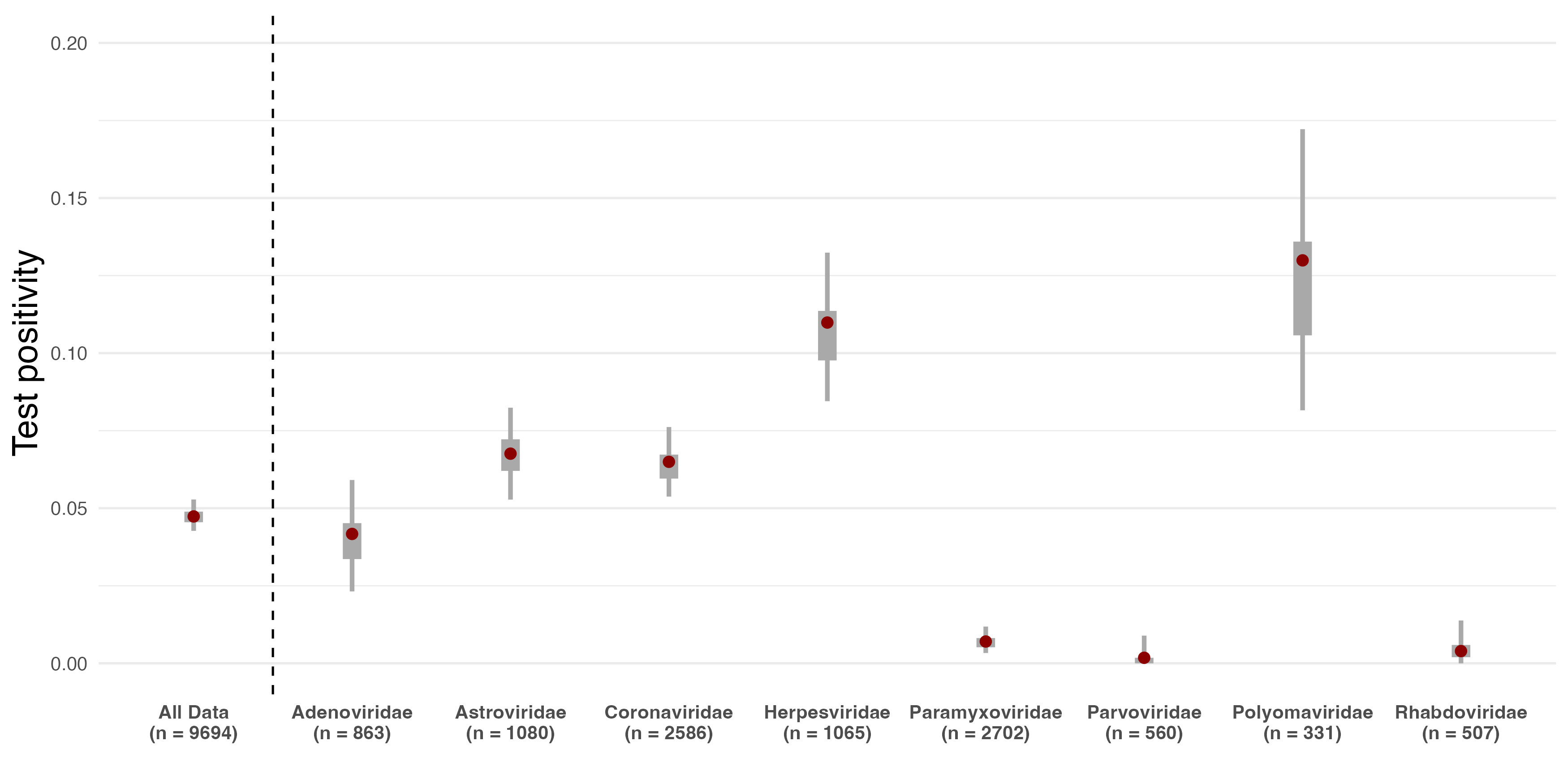

**Figure S6. Ridgeline plots showing species-specific intercepts (a), pregnancy effects (b), and lactation effects (c) from a hierarchical Bayesian model of viral detection in female bats.** Ridgeline plots show the full posterior distribution for every varying effect parameter. All parameter estimates are presented on the log-odds scale. In each plot, the posterior distributions have been sorted according to their median values. Detailed description of the model structure, predictor variables, and model fitting is given in the main text.

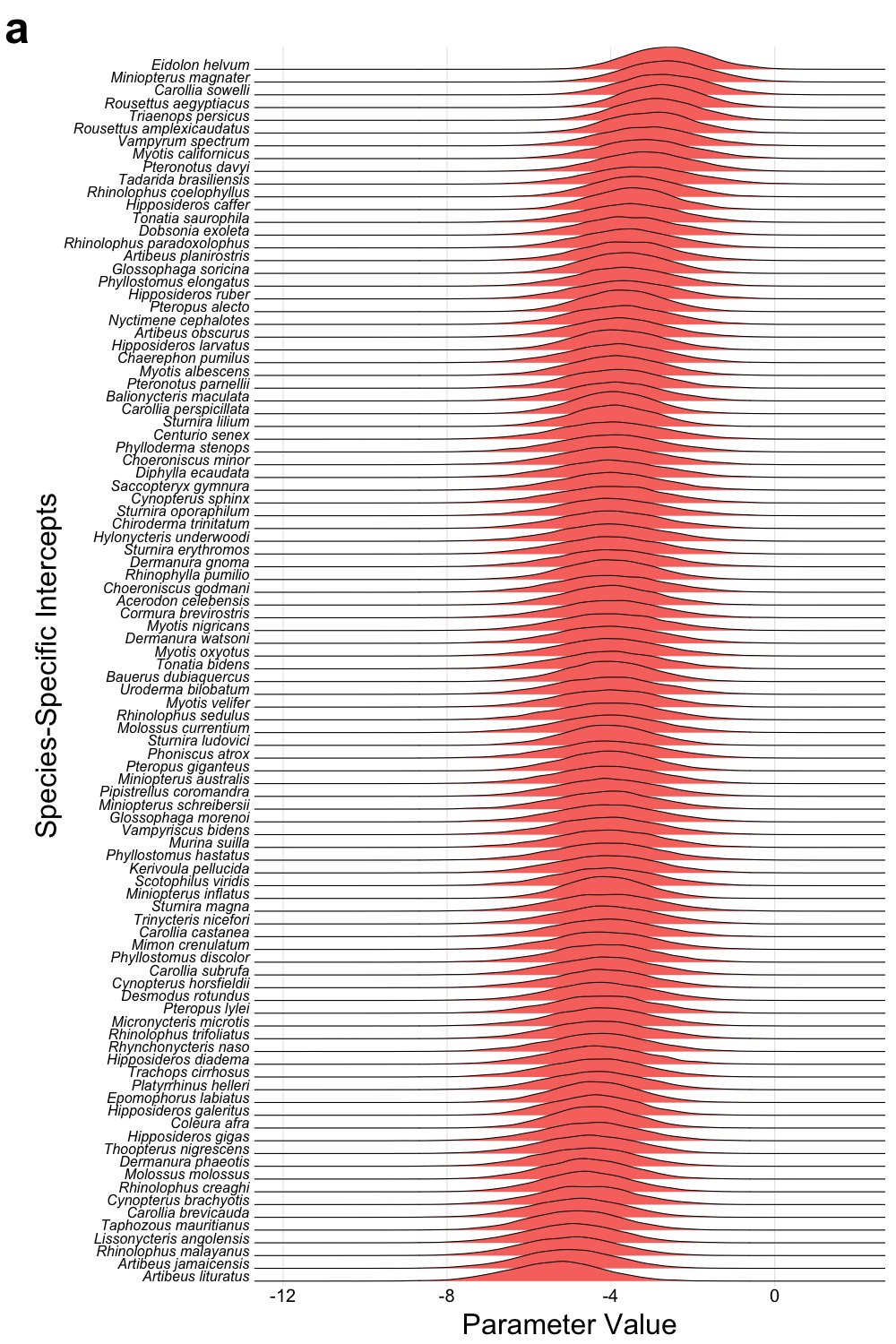

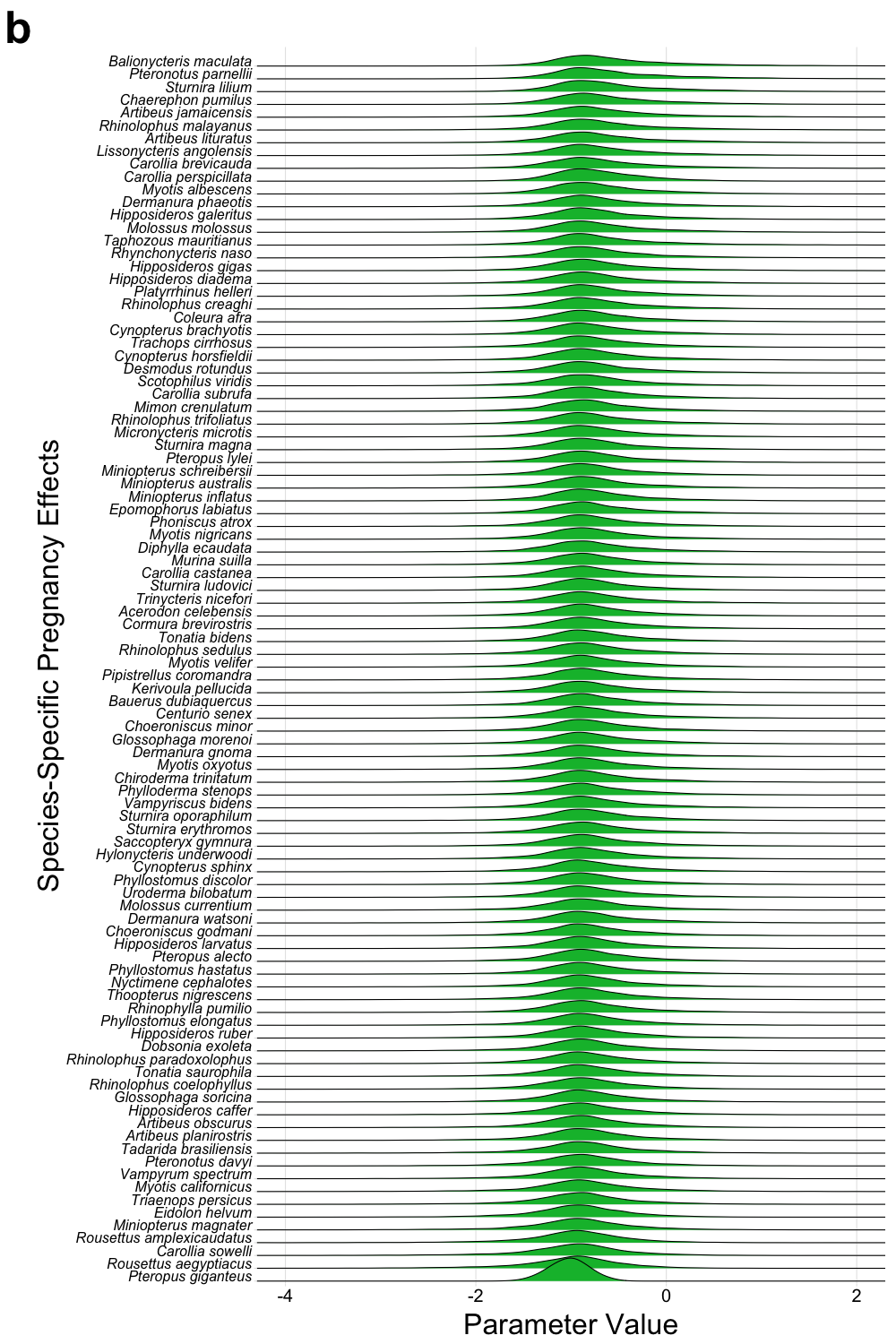

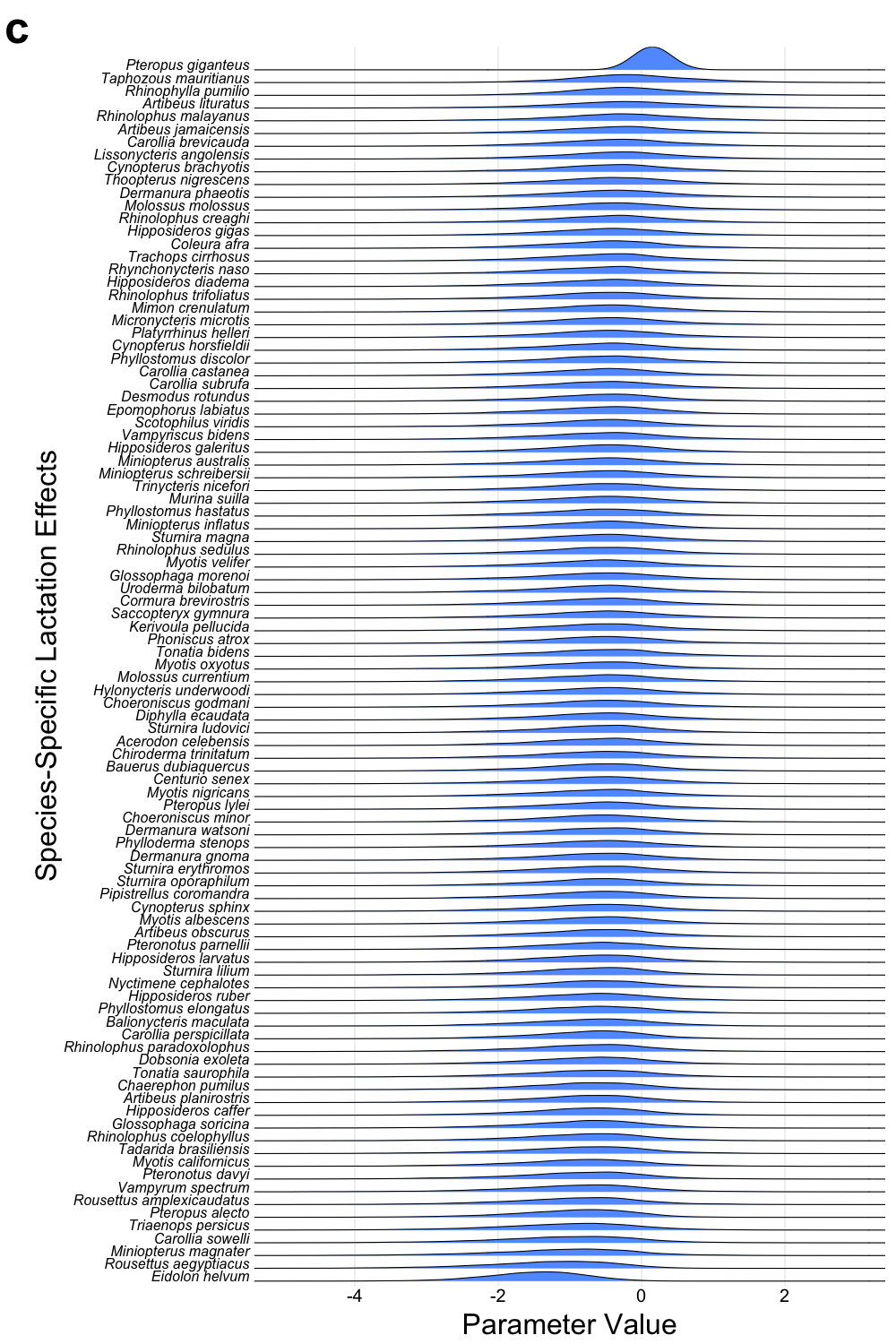

**Figure S7. Ridgeline plots showing all varying intercepts clusters from a hierarchical Bayesian model of viral detection in female bats.** Ridgeline plots show the full posterior distribution for every varying intercept parameter across the five varying intercepts clusters in the model. All parameter estimates are presented on the log-odds scale. In (a), intercept labels have been excluded to simplify presentation and encourage visual comparison of relative variation among the clusters. Labels indicating the intercept estimate identities appear in (b-f). X-axis limits have also been modified as appropriate in (b-f), however, posterior distributions are identical to (a). In each plot, the posterior distributions have been sorted according to their median values. Detailed description of the model structure, predictor variables, and model fitting is given in the main text.

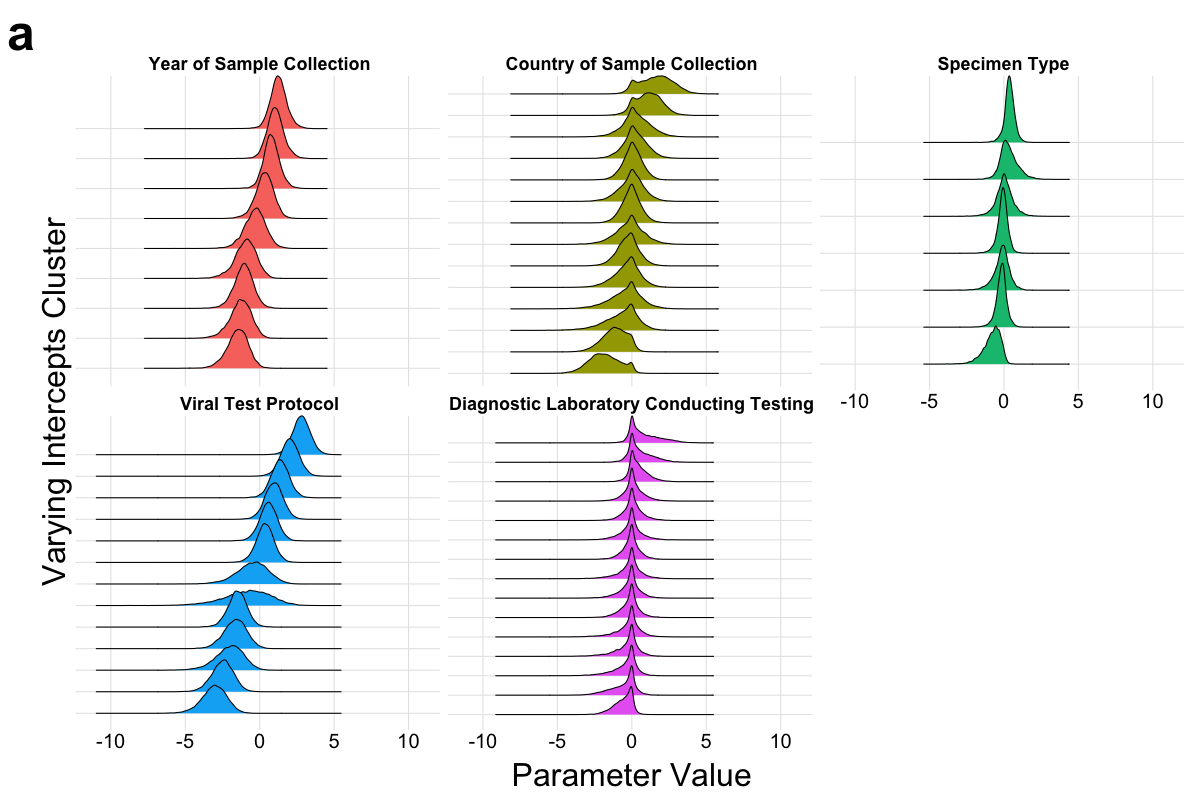

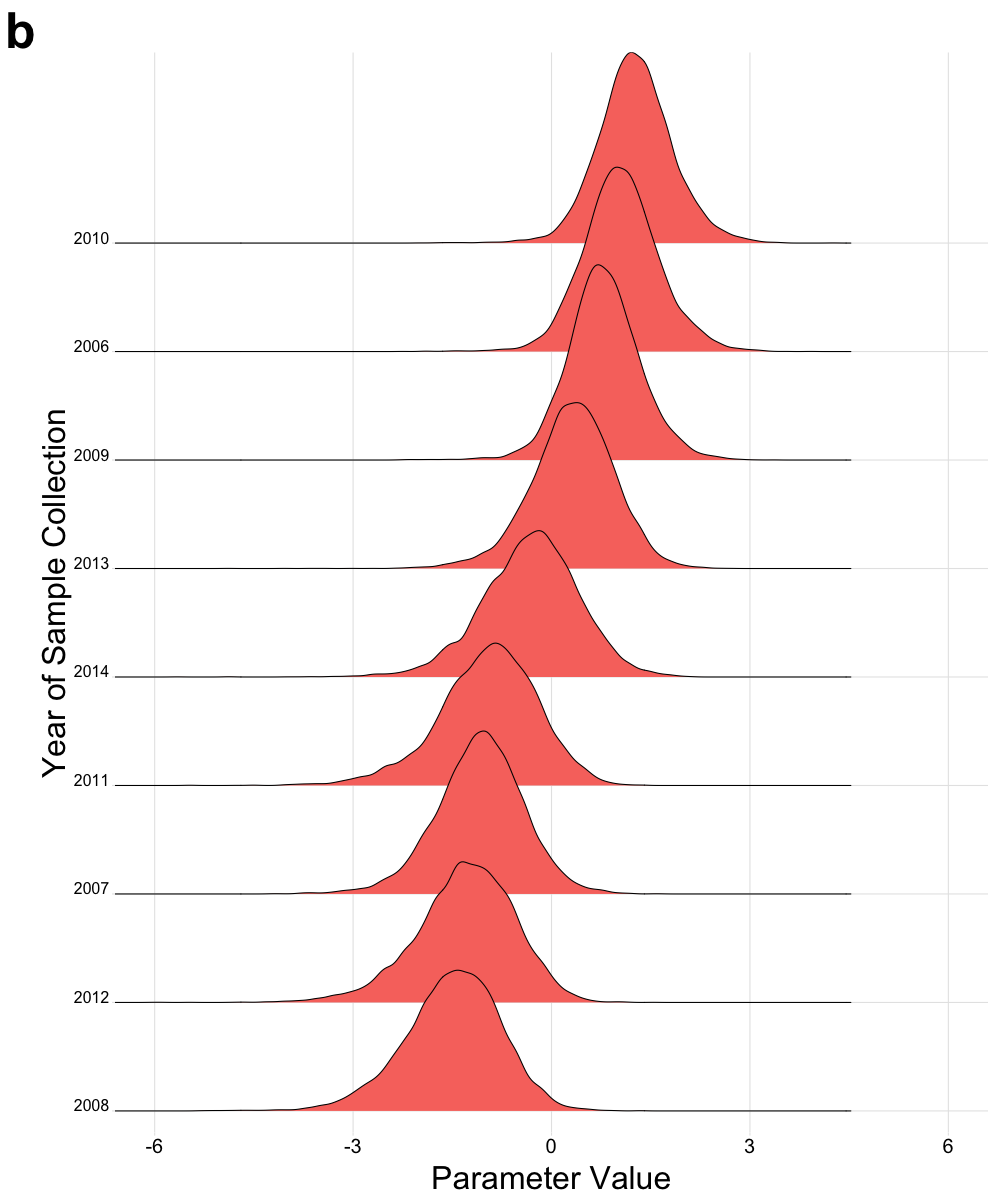

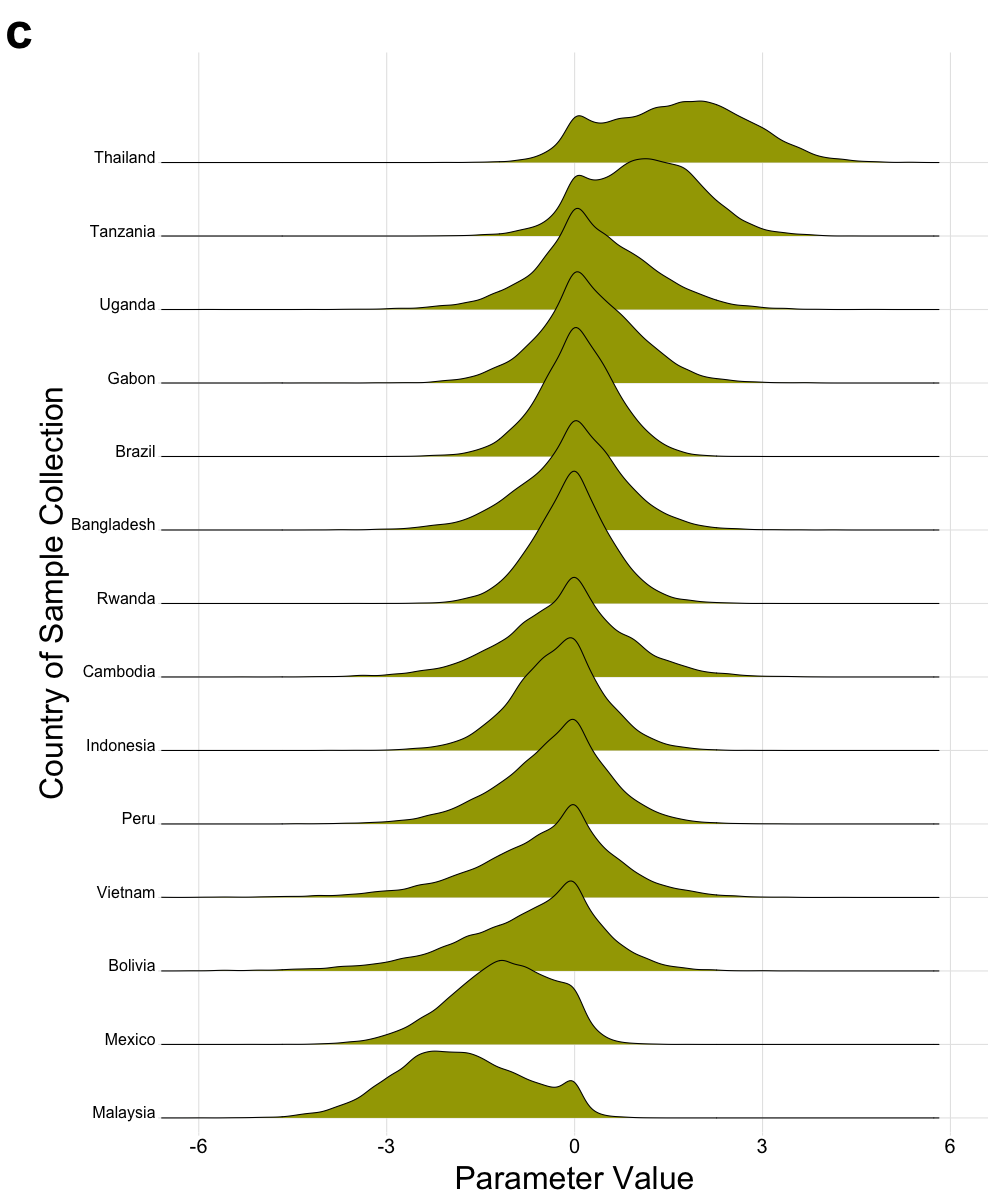

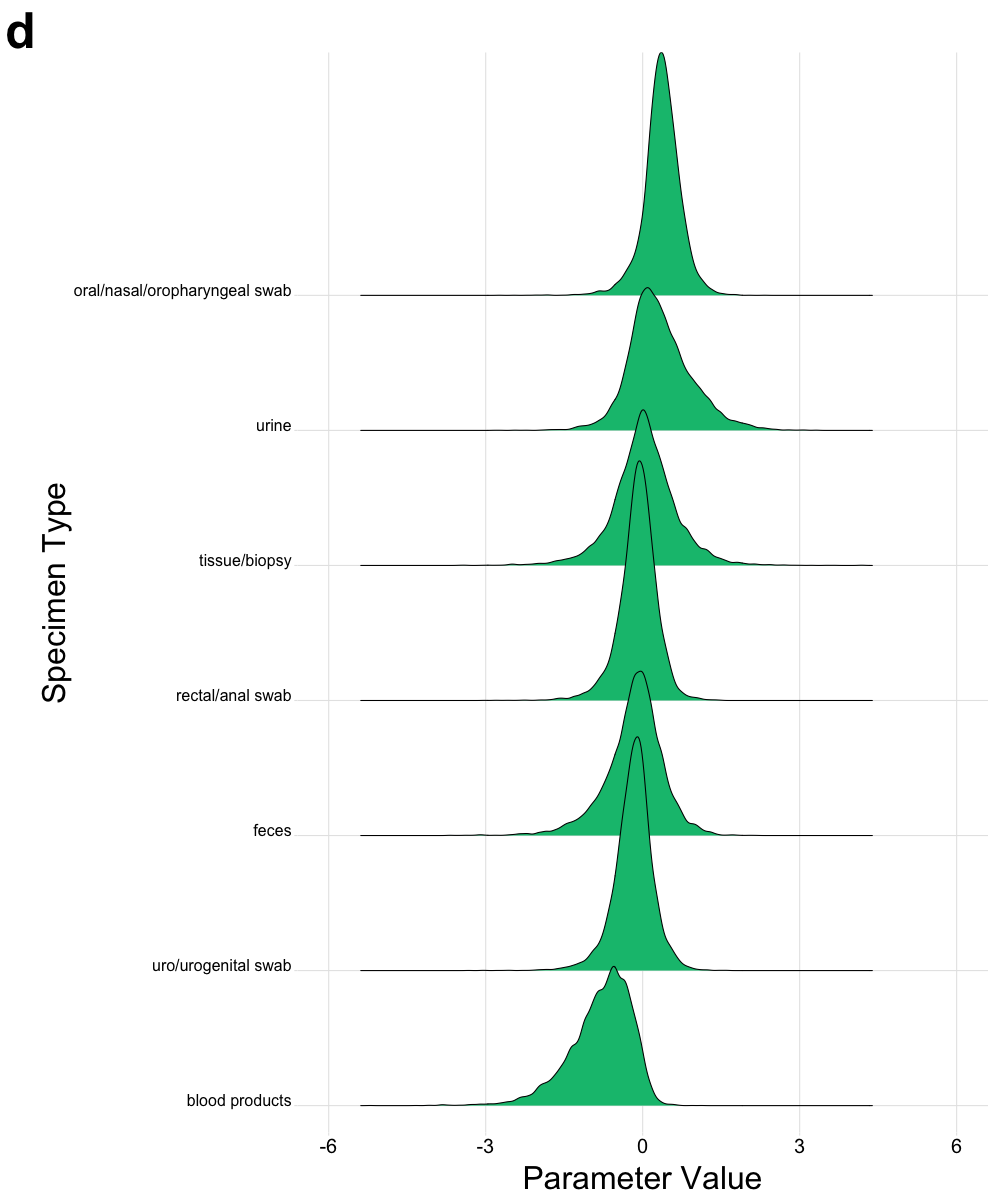

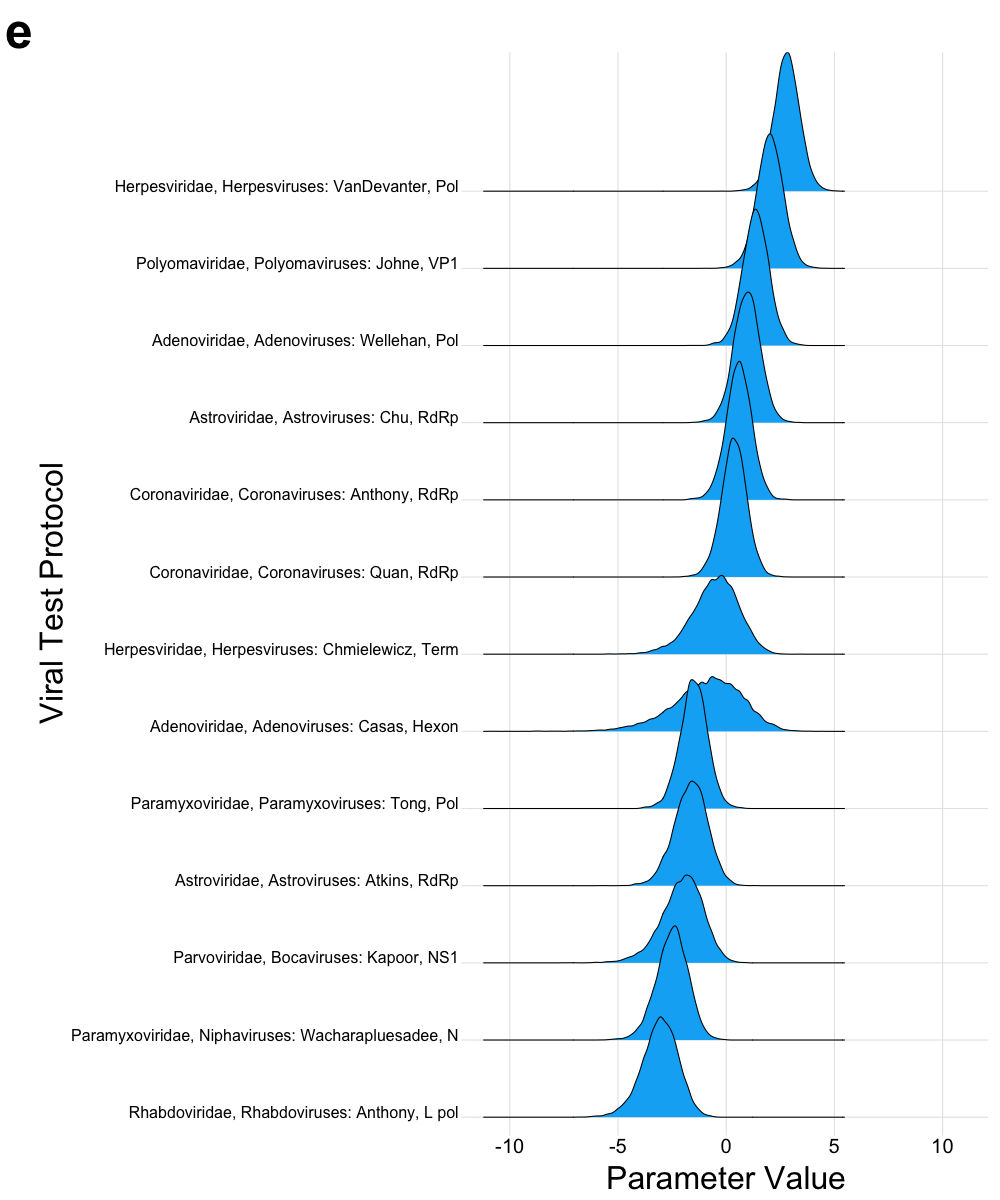

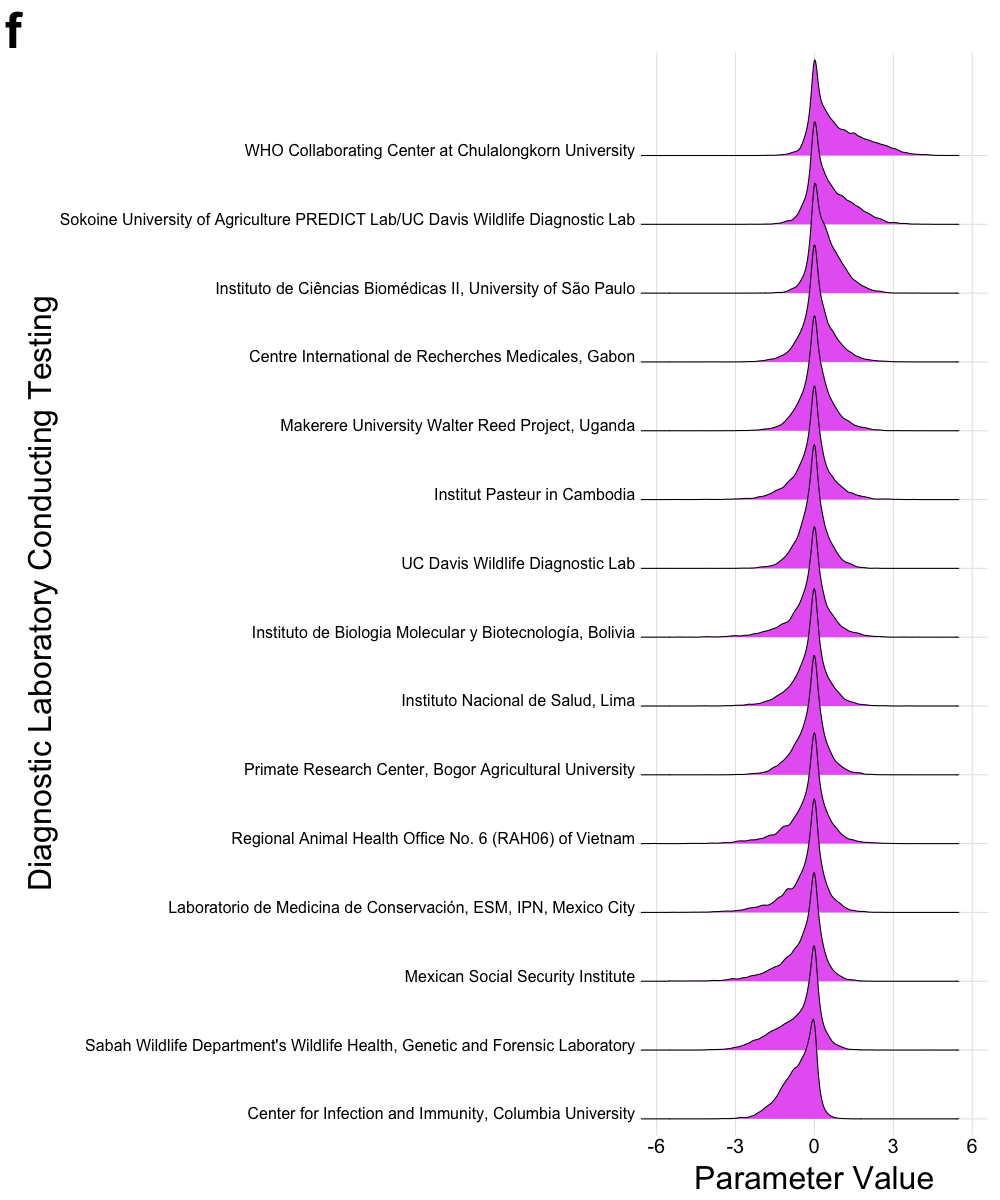
